# Water parameters and hydrodynamics in rivers and caves hosting *Astyanax mexicanus* populations reveal macro-, meso- and micro-habitat characteristics

**DOI:** 10.1101/2025.02.06.635682

**Authors:** Laurent Legendre, Stéphane Père, François Rebaudo, Luis Espinasa, Joël Attia, Sylvie Rétaux

**Author notes:** Authors for correspondence &.

## Abstract

The Mexican tetra (*Astyanax mexicanus*) has emerged as a leading model for evolutionary biology and the study of adaptation to extreme subterranean environments. The river-dwelling morph of the species is distributed in Mexico and Texas, while the blind and cave-adapted morph inhabits the karstic caves of the Sierra Madre Oriental in northeastern Mexico. The molecular, cellular and genetic underpinnings of *Astyanax* cavefish evolution are increasingly studied, but our understanding of its habitat and environment is incomplete, limiting the interpretations of its morphological, physiological, and behavioral adaptations. In particular, the physico-chemical parameters of the water and the hydrological regimes to which cavefish are subjected are largely unexplored. From 2009 to 2025, we have recorded the physico-chemical parameters of the water at localities hosting *Astyanax mexicanus* cavefish and surface fish in the Sierra de El Abra and Sierra La Colmena regions of the states of San Luis Potosí and Tamaulipas, Mexico. We sampled 13 caves out of the 33 known *Astyanax* caves and 30 surface stations (rivers, springs, ponds). Data were collected using a variety of devices and probes, including both point measurements (at the end of winter) and longitudinal measurements (throughout the year). The comparison of epigean and hypogean waters showed strong signatures of the two macro-habitats. As compared to surface, on average cave water was cooler, much less conductive and highly anoxic. Moreover, a comparison between different caves (i.e., meso-habitat level) revealed significant differences in both specific water parameters and hydrological regimes. One- or two-year longitudinal recordings demonstrated that some caves exhibit relatively stable hydrological regimes, while others experience multiple, sudden and significant fluctuations. Finally, distinct pools within a single cave showed notable differences, displaying a reproducible increasing gradient in water temperature as a function of distance from the cave entrance, and revealing specificities at the micro-habitat level. We interpret our comprehensive dataset on cave water quality and hydrodynamics in the context of an integrated view of cave biology and the evolution of cave organisms.

## Introduction

The environmental parameters of an ecosystem are crucial for defining a species’ ecological niche. The range of these parameters within the species’ habitat helps to identify the preferred conditions for that species. In aquatic environments, water serves as the medium for all organisms, making the study of its physico-chemical properties essential for understanding the species’ parameter range and, consequently, their biology, eg (Reichard, 2016). Comprehending these parameter ranges is key to understanding the species’ adaptive responses, evolutionary history, and distribution within its ecological niche. These factors are also essential for conservation efforts and provide the foundation for managing the species’ health, welfare, care and husbandry in captivity. This is particularly relevant when the species is used as an animal model in laboratory settings, where reproducible experiments are required across a range of disciplines, including biomedical research.

The fish *Astyanax mexicanus*, or Mexican tetra (Sardinita Mexicana and Sardinita ciega), has become an excellent model for Eco-Evo-Devo, a concept first coined by Gilbert in 2015 (Gilbert et al., 2015). This is because this single species exists in two forms, or eco-morphs: a freshwater river-dwelling morph that is widely distributed in Mexico and the southern United States; and a blind, depigmented dark-adapted morphs that inhabits the 33 caves discovered so far in a restricted area of northeastern Mexico (Elliott, 2018; Espinasa et al., 2018; Espinasa et al., 2020; Jeffery, 2001; Jeffery, 2008; Jeffery, 2009; Miranda-Gamboa et al., 2023; Mitchell et al., 1977). Of the more than 300 currently known species of cave and groundwater fishes (Proudlove), *Astyanax mexicanus* is by far the most studied (Rétaux and Jeffery, 2023). The availability of the two lineages, surface and cave, offers unique opportunities for evolutionary biology and genetics (Casane and Rétaux, 2016; Gross et al., 2023; Gross et al., 2009; Protas et al., 2008; Protas et al., 2006). In the last twenty years, much has been discovered about the neurosensory, behavioral and physiological adaptations of *Astyanax* cavefish, e.g. (Aspiras et al., 2015; Blin et al., 2024; Hinaux et al., 2016; Riddle et al., 2018; Yoshizawa et al., 2010; Yoshizawa et al., 2014), and progress has been made in understanding the evolution of its genome (Krishnan et al., 2022; Leclercq et al., 2024; McGaugh et al., 2014; Warren et al., 2021). However, population biology studies have sometimes yielded conflicting results on the origin, structure, and age of cavefish populations (Bradic et al., 2012; Fumey et al., 2018; Herman et al., 2018; Policarpo et al., 2021; Policarpo et al., 2024; Strecker et al., 2004) and laboratory and field studies are sometimes difficult to reconcile [e.g. (Blin et al., 2024) vs. (Blin et al., 2020); (Aspiras et al., 2015) vs. (Espinasa et al., 2017; Simon et al., 2017)].

The necessity of a comprehensive understanding of troglomorphic evolution in cavefish, in response to the profound environmental changes associated with cave life, has been highlighted (Rétaux and Jeffery, 2023; Torres-Paz et al., 2018). This includes a better understanding of the ecosystems in which *Astyanax* live, starting with long-term monitoring and comparison of surface and cave water quality. However, there is a “habitat impediment” related to the objective difficulties of exploring subterranean habitats (Mammola et al., 2021). To our knowledge, very few studies have longitudinally documented water quality in caves in relation to cavefish biology - except to discuss the variation of condition factor across seasons for *Ituglanis* sp. in Brazilian caves (Bichuette and Trajano, 2021), or to discuss the evolution of temperature preference in some *Astyanax* cavefish populations (Tabin et al., 2018). The hydrological regimes are also largely unknown. The 2004 report by Fish (corresponding to his PhD thesis in 1976) describes the karst hydrogeology and geomorphology of the Sierra de El Abra region, which includes some of the *Astyanax* caves, and provides some temperature and pH data from surface and cave stations, but the work is disconnected from the biological perspective of the *Astyanax* cavefish (Fish, 2004). To fill this gap, here we present the results of 15 years of efforts to record physicochemical parameters of the water in several cave and surface localities where *Astyanax mexicanus* fishes swim, with both point and longitudinal measurements. We uncover large, sometimes unexpected differences in water quality and hydrodynamics between surface streams and caves (macro-habitats), between caves (meso-habitats), and between pools within a cave (micro-habitats). We then discuss our results in terms of the evolution and biology of *Astyanax* Mexican cavefish.

## Results

Between 2009 and 2025, we have systematically recorded the physico-chemical parameters of the water in which *Astyanax mexicanus* cavefish and surface fish swim. We collected data using several devices and probes (see Methods), both with point measurements (“one-off”, mostly at the end of winter) and longitudinal measurements of water quality. We sampled 13 caves out of the 33 known *Astyanax* caves and 30 surface stations (rivers, springs, ponds), mainly in the Sierra de El Abra and Sierra La Colmena regions of the states of San Luis Potosi and Tamaulipas, Mexico (Fig. 1AB, Fig1-figure supplement 1 and Supplemental File1).

**Figure 1:**
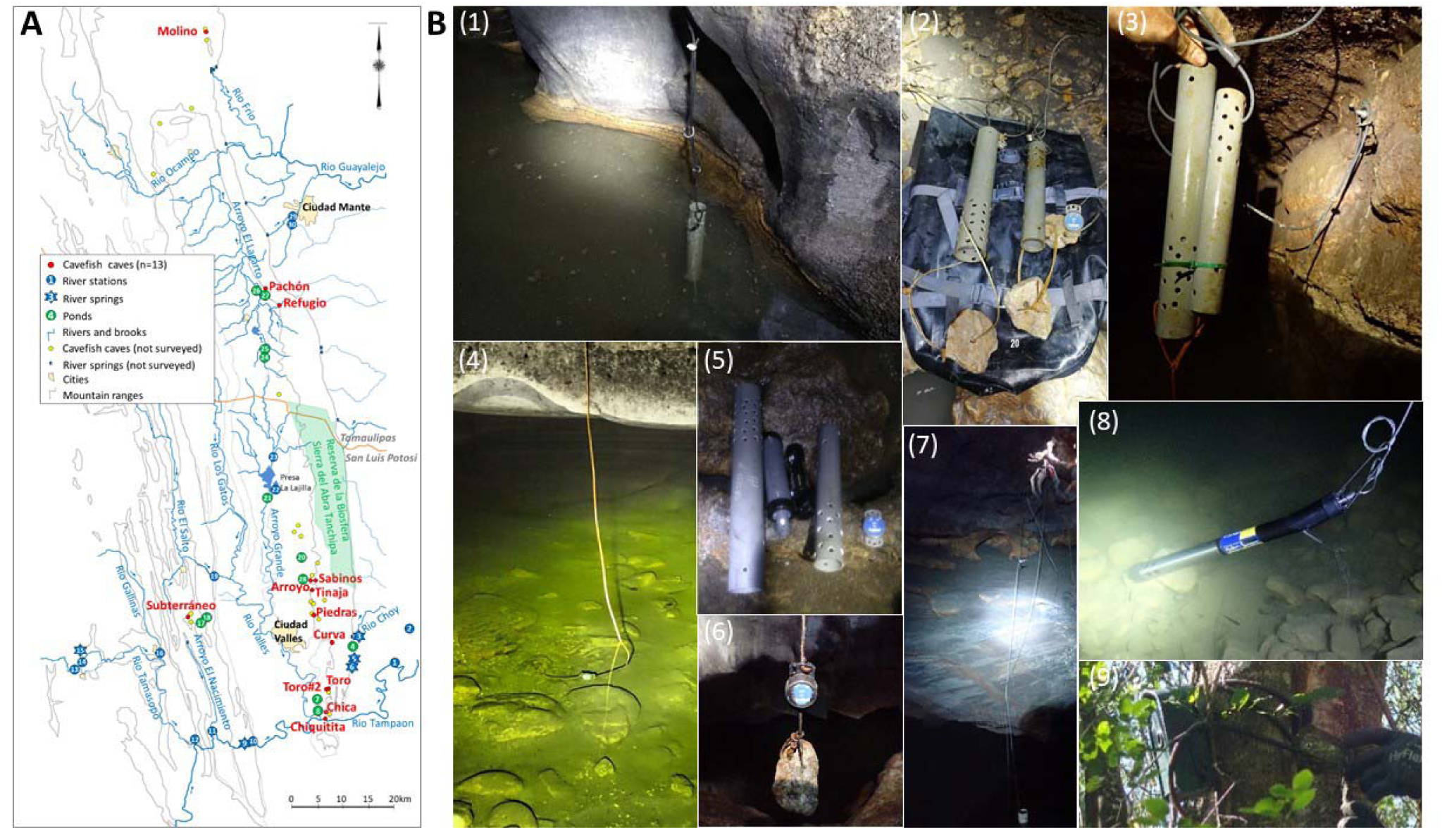
Water sampling stations and equipment used. A: Map of the area. See inset for symbols and legends. B: Examples of probes used and their installation on sites. (1) Hobo probes (temperature, pH, conductivity) protected in plastic tubes in Chica pool 2.5. (2) Recovery of probes and their protection and attachment devices in Subterráneo. (3) Fixation and securing of two probes in Sabinos pool1. (4) A small Hobo probe installed in shallow water in Sabinos pool2. (5) Preparation of pH and conductivity probes (Hobo) and their protection tubes in Chica. (6) State of conservation of a small Hobo probe after 3 years (2019-2022) in Sabinos pool1. (7) Fixation and securing of the multiparametric probe (In-Situ) in the back of Pachón main pool. (8) Point measurement with multiparametric probe in Subterráneo pool2. (9) Fixation and securing of camera trap for time-lapse recording of water regime at the entrance of Chica. All photos by LL and SR. See also Supplemental Figure 1.

### Comparing physico-chemical parameters of the water at the surface and in underground caves by point measurements at the end of winter

Over the years, we have carried out 183 point measurements in caves and 77 point measurements in surface waters for temperature, pH and Electrical Conductivity (EC, abbreviated “conductivity” below)(Fig. 2A and Table 1, first two lines). These point measures were taken in February-March and therefore represent the situation at the end of winter/end of the dry season. From this dataset, the temperature in surface waters was on average 1.24°C higher than in caves (mean surface 24.86°C; mean cave 23.59°C; t = -3.8, df = 101.8, p-value = 1.8e-5), ranging from 19.3°C to 31.1°C at the surface and from 16.8°C (Sótano del Arroyo, 21 February 2024) to 27.5°C (Cueva Chica, 19 February 2024) in caves. The pH was very similar in the two environments (mean 7.67 in caves vs 7.78 in surface waters; t = -1.3, df = 98.3, p-value = 0.18). The conductivity was twice higher in surface water (mean 1092 µS.cm) than in cave water (mean 508 µS.cm; t = -10.7, df = 68.1, p-value = 2.5e-20). Therefore, epigean and hypogean waters seem globally different, especially in terms of conductivity and, to a lesser extent, temperature. Principal Component Analysis (PCA) followed by an ANOVA on the first PCA axis (pca1) further demonstrated that surface and cave waters display a strong physico-chemical signature, with 95% confidence ellipses around centroids showing no overlap, with temperature and conductivity having a major contribution to the segregation of the two clusters in the first dimension (ANOVA on pca1, F = 27.597, P = 6.29e-07) (Fig. 2C, Fig2-figure supplement 1).

**Figure 2:**
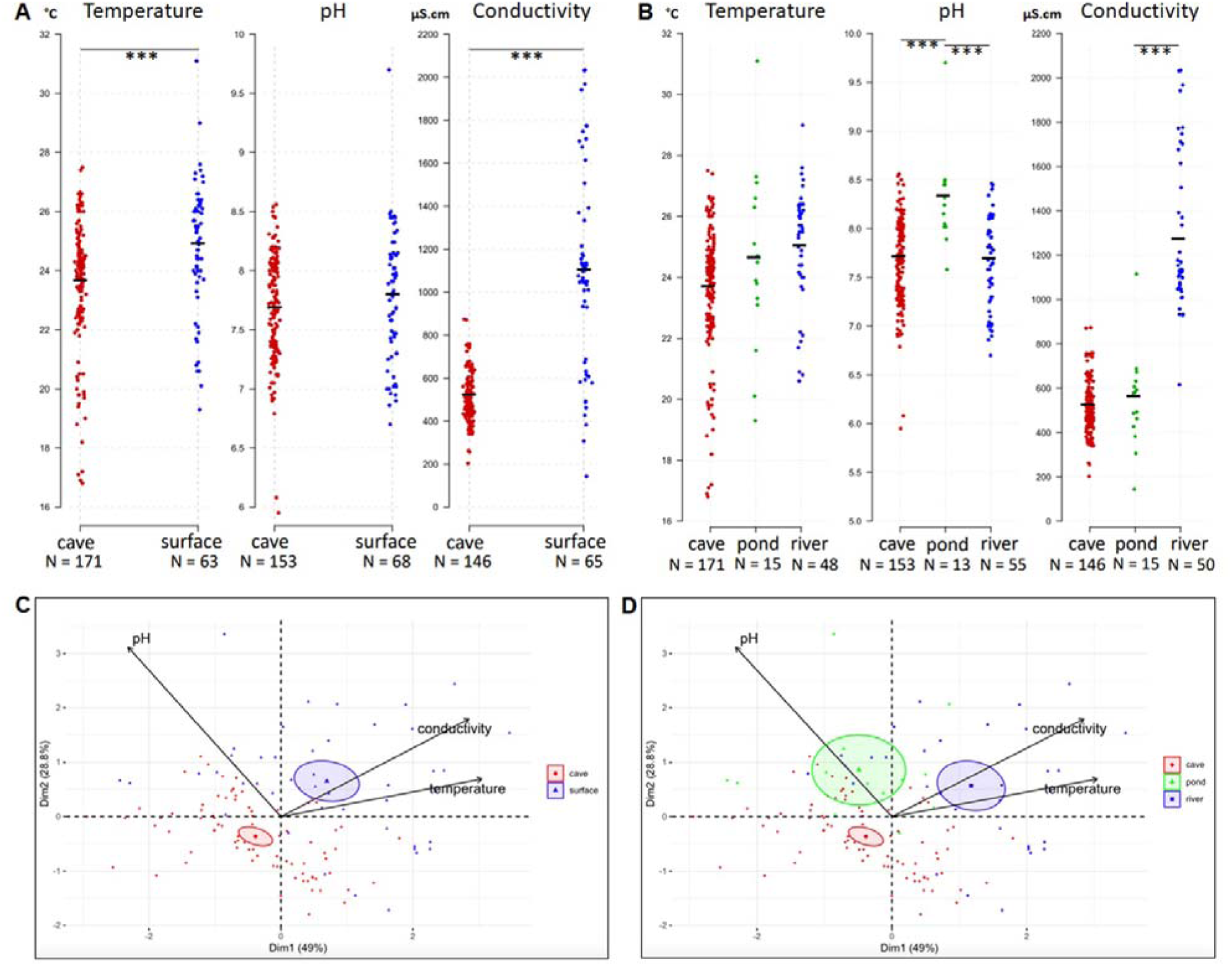
Comparing physico-chemical parameters of the water from point measurements in surface waters and in underground caves (February-March measurements). A, B: distribution and comparison of temperature, pH and conductivity between cave (red dots) and surface waters (blue dots) (A) and between cave waters (red dots), still surface water (green dots), and running surface waters (blue dots) (B). Wilcoxon - Mann Whitney tests (see Table1). C, D: Clustering of the physico-chemical properties of the water using PCA. Color codes are indicated, 95% confidence ellipses are shown. This graph includes values recorded with Combo Hanna. Ncaves=81, Nsurface=45, N=3 variables. See Supplemental File 2 (raw data) and Fig.2-figure Supplement1. Color codes are indicated, 95% confidence ellipses are shown. C, ANOVA on pca1, F = 27.597, P = 6.29e-07; ANOVA on pca2, F = 46.636, P = 3.39e-10. D, ANOVA on pca1, F = 29.003, P = 4.79e-11; ANOVA on pca2, F = 23.965, P = 1.62e-9.

**Table 1:** Summary statistics for point measurements of water physico-chemical parameters in caves, and in still and running surface waters (2009-2024; February-March measurements). a, comparison between cave and surface. b, comparison between running and still surface water. Wilcoxon - Mann Whitney tests.

|  | Sampling dates | Temperature (°C) |  |  |  | pH |  |  |  | Conductivity (µS.cm) |  |  |  |
| --- | --- | --- | --- | --- | --- | --- | --- | --- | --- | --- | --- | --- | --- |
|  | 2009-2024 | mean | median | min | max | mean | median | min | max | mean | median | min | max |
| Caves, all | 2009-2024 | 23.59<br>N=171 | 24.06 | 16.8 | 27.5 | 7.67<br>N=153 | 7.69 | 5.95 | 8.56 | 508<br>N=146 | 490 | 203 | 874 |
| Surface, all | 2009-2024 | 24.86<br>N=64 | 25.3 | 19.3 | 31.1 | 7.78<br>N=68 | 7.82 | 6.7 | 9.7 | 1092<br>N=65 | 1098 | 144 | 2034 |
| a |  | p=1.8e-5 |  |  |  | p=0.18 (NS) |  |  |  | p=2.5e-20 |  |  |  |
| Surface, running | 2009-2024 | 24.92<br>N=48 | 25.5 | 20.6 | 31.1 | 7.65<br>N=55 | 7.66 | 6.7 | 8.46 | 1256<br>N=50 | 1127 | 616 | 2034 |
| Surface, still | 2009-2024 | 24.66<br>N=15 | 24.5 | 19.3 | 29 | 8.32<br>N=13 | 8.24 | 7.58 | 9.7 | 545<br>N=15 | 578 | 144 | 1116 |
| b |  | p=0.68 (NS) |  |  |  | p=0.00019 |  |  |  | p=5.2e-8 |  |  |  |

Despite significant differences emerging between the two environments, we noticed a large dispersion of the values in our sampling (Fig. 2A). For example, conductivity in surface waters varied from 144 µS.cm (station #4 on map in Fig. 1A; small pond near Camino Taninul, 2013) to 2034 µS.cm (station #15; Arroyo Los Otates, spring/resurgence close to la Cueva del Fraile, 2019), i.e., a 14-fold variation. Similarly, station #4 displayed an exceptionally high pH value of 9.7 – but despite this extreme condition, a number of surface *Astyanax* swam there together with many black tadpoles of an undetermined anuran species. We reasoned that such data dispersion might be due to different conditions encountered at different surface locations, and we replotted the data after sorting the sampling sites into “running” and “still” water. The term "running" includes here any river station with flowing water as a typical lotic ecosystem. The term "still" includes any other water unit such as a pond (including a pond in a river during the dry season), a lake, or a well, as a typical lentic ecosystem (Fig. 2B and Table 1, last two lines). The mean temperature was similar in still and running water at the surface (24.92°C vs 24.66°C), and higher than in caves (23.59°C). The mean pH was higher in still water than in running water (8.24 vs 7.66; p=0.00019). In addition, caves and flowing surface rivers displayed the same mean pH (7.64 and 7.65, respectively). Conversely, regarding conductivity still water bodies were similar to caves (545 µS.cm and 508 µS.cm) and both were significantly different from running water (surface running 1256 µS.cm; surface still 578 µS.cm; caves 508 µS.cm; p=5.2e-8). A PCA confirmed that surface running, surface still and cave waters segregate well in the physico-chemical space, suggesting that each of these three environments has its own signature (ANOVA on pca1, F = 29.003, P = 4.79e-11; ANOVA on pca2, F = 23.965, P = 1.62e-9) (Fig.2D). Complementary Tukey contrast tests revealed a significant difference between river and pond/cave on pca1 values (river *vs* cave: t = -7.389, P < 0.0001; river *vs* pond: t = -5.006, P < 0.0001) and between cave and pond/river on pca2 axis (cave vs pond: t = -5.115, P < 0.0001; cave vs river: t = -5.585, P < 0.0001). In summary, the caves, the ponds and the rivers of the sampled area have different combinations of physico-chemical water parameters, and some extreme values can be observed. All the waterbodies we sampled hosted *Astyanax mexicanus* fish (surface- or cave-dwelling morphs), suggesting that the species is adapted to rather diverse ecological conditions (see also below).

### Stability of measurements over time (1953-2025)

We sought to test whether the parameters analyzed might have varied over time. We compared our data (2009-2025) to earlier measures reported in bibliography (Bonet, 1953; Elliott, 2018; Fish, 2004; Mitchell et al., 1977; Ornelas-Garcia et al., 2018; Tabin et al., 2018); Johnson, 1967). One hundred and fifteen point measures were found between 1953 and 2022, in 8 different papers, mostly regarding temperature. A majority of these measures were found in Fish 1977 and 2004 (n=65), Tabin 2018 (n=17), Elliott 2018 (n=14) and Johnson 1967 (n=8). For our comparison, we chose one cave (Pachón) and one surface site (Nacimiento del Río Choy; station #3 on map Fig. 1A), i.e. the stations where the most repeated samples are available over the years and where the exact sampling location is unambiguous. For conductivity, there was no data in the literature. For pH, measures by Fish in 1977 were similar to ours - but note data from Fish are mostly from June and July (beginning of rainy season) whereas our data are from February-March (dry season, end of winter). Regarding temperature, the old measurements showed values in the same range as our recent ones or some other recent measures by other authors, both for Pachón cave and for the Nacimiento del Río Choy spring (Fig. 3, see legend), suggesting that temperatures in these two sites have remained stable over 7 decades. The sparser data available from 4 other caves show the same trend (Fig3-figure supplement 1, see also legend).

**Figure 3:**
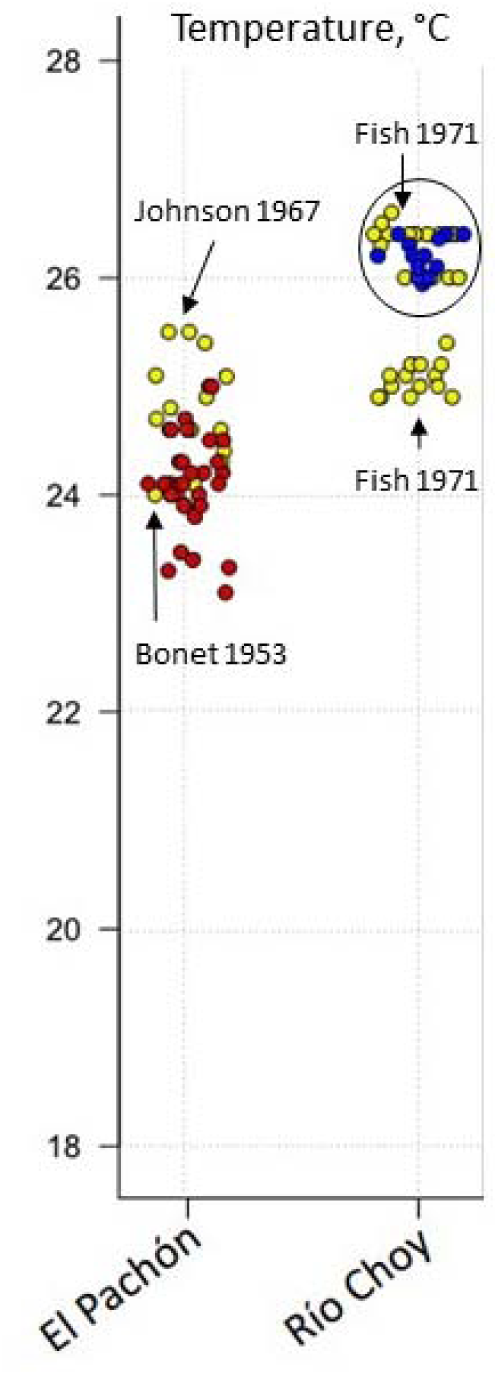
Stability of water temperature measurements over 7 decades (1953-2024) Red dots (Pachón cave) and blue dots (river, Nacimiento del Río Choy) are from our dataset. Yellow dots are from ancient literature, with author’s name & date. For the surface station Nacimiento del Río Choy (blue dots; river location #3 on Fig.1A), a black circle shows a group of ours & old data (Fish, 2004) close to 26°C. Another group of only older data is close to 25°C, also from Fish (2004). Fish showed that temperature could vary within days with flooding (e.g., between 5 June and 14 August 1971) at this location, with also massive variations in discharge, turbidity and water chemistry.

### Comparing physico-chemical parameters of the water between different caves, and between different pools in a single cave

Recent field studies have shown that the biology and behavior of cavefish can vary significantly between caves and between pools within a cave (Blin et al., 2020; Espinasa et al., 2021; Hyacinthe et al., 2023; Simon et al., 2017), suggesting that cavefish may adapt to local environmental conditions. Since water is the most important environmental variable, we hypothesized that temperature, pH and conductivity can vary significantly between caves and within caves. Below, we compared data recorded in several pools of 6 different caves (Fig. 1A and Supplemental File1): Cueva de El Pachón, Cueva de Los Sabinos, Sótano de la Tinaja, Sótano de Las Piedras, Cueva Chica (El Abra caves), and Cueva del Río Subterráneo (Micos area) (Fig. 4).

**Figure 4:**
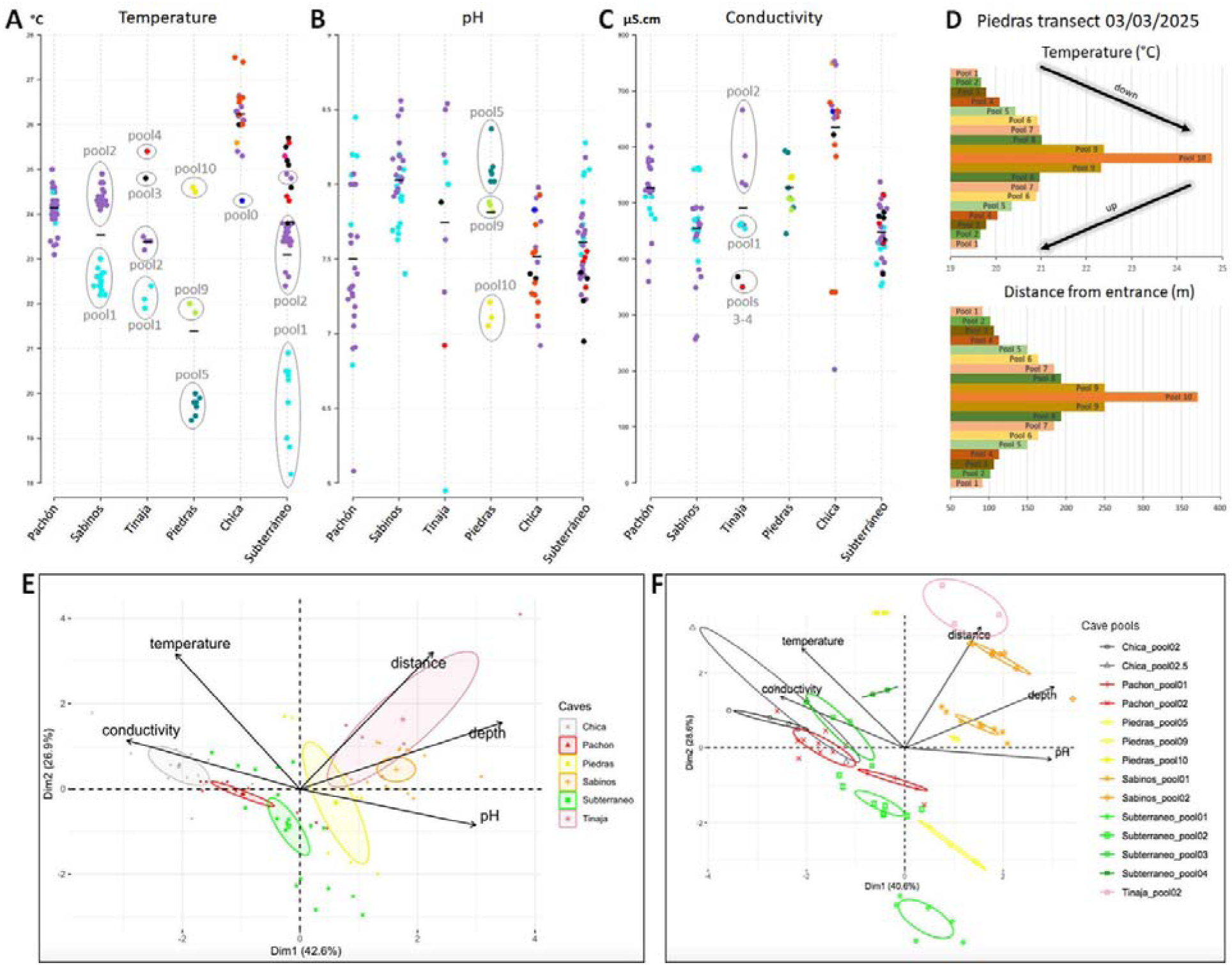
Comparing physico-chemical parameters of the water between different caves, and between different pools in a single cave (February-March measurements) A: Temperature. B: pH. C: Conductivity In the 3 graphs A-B-C, in each cave, dots are color-coded according to the pool in which measurements were made. When useful, pool numbers are indicated, with number 1 being closest to the cave entrance, and higher numbers corresponding to successive pools encountered, further away from entrance and/or deeper in the cave. D: Transect analysis in Piedras cave through pools 1 to 10 (03/03/2025), showing temperature and distance from entrance. The low-to-high temperature gradient from the first to last pool encountered seems to vary as a function of the distance from entrance. E, F: Clustering of the physico-chemical and geomorphological properties of caves and cave pools using PCA. Color codes are indicated, 95% confidence ellipses are shown. E, ANOVA on pca1, F = 48.920, P < 2.2e-16; ANOVA on pca2, F = 7.899, P = 7.04e-6 F, (ANOVA on pca1, F = 25.543, P < 2.2e-16; ANOVA on pca2, F = 68.433, P < 2.2e-16

Temperatures were different between caves when considering the entire dataset (mean water temperature in Chica: 26.19°C; Pachón: 24.1°C; Sabinos: 23.49°C; Tinaja: 23.35°C; Subterráneo: 23.05°C; Piedras: 21.35°C; maximum variation: 4.84/18%), with clear tendencies for Chica to be a “hot cave” and for Piedras to be a “cool cave” (Fig. 4A). Moreover, temperatures were different between different pools in a given cave (Fig. 4A). Indeed, at the end of winter (February-March measures) in all caves the water temperature increased as a function of the depth and/or the distance of the pools from cave entrance (see cave maps in Supplemental File1). In all cases, pools located close to cave entrances were cooler than deeper/more distant pools close to final sumps, with a remarkable gradient of water temperature. This pattern applies globally to all studied caves except Pachón: this cave is small as compared to others, and the so-called main pool and lateral pool are a few meters apart and form a single body of water that communicates most of the year when water level is high, therefore recording the same temperature in the two pools was expected. Moreover, the pools in the Maryland passage are almost never accessible for measures (Supplemental File1 and see below). Conversely, in Piedras for example, pool5 (the first large pool encountered at ∼150m from pit entrance, at a ∼30m depth) has an average temperature of 19.7°C, while pool9 (250m from entrance, 45m deep) has an average temperature of 21.9°C and the final pool10 (370m from entrance, 47m deep) has a temperature of 24.6°C. Therefore, there is a 4.9°C difference between these pools of the same Piedras cave, all of which host cavefish (Fig. 4A). To understand further this pattern, in March 2025 we performed systematic transect measurements in the ten successive pools of Piedras (Fig. 4D and Fig4-figure supplement 1, Supplemental File1). We confirmed the water temperature gradient, which was progressive and pronounced, and well related to the distance of the pool from cave entrance (but not to depth, Fig4-figure supplement 1). Thus, the combined effects of the outside air (fresh air in the entrance pit covered by rain forest in the case of Piedras), of the cave topography, and of the presence of a large bat colony in the distal part of the cave seem to contribute to the establishment of such notable temperature gradient across the different cave pools.

Average pH values were similar in all studied caves (Sabinos: 8.01; Piedras: 7.8; Tinaja: 7.7; Subterráneo: 7.58; Chica: 7.48; Pachón: 7.4; maximum variation: 0.61/7%), with Sabinos showing the most basic pH values. There was no such pattern as seen with temperature between pools within a cave, as confirmed by the systematic transect study in Piedras (Fig4-figure supplement 1).

Finally, average conductivity was also variable between caves (Chica: 631 µS.cm; Piedras: 525 µS.cm; Pachón: 524 µS.cm; Tinaja: 487 µS.cm; Sabinos: 451 µS.cm; Subterráneo: 444 µS.cm; maximum variation: 187/29%), as well as between pools within a cave. For example in Tinaja conductivity values were comprised between 359µS.cm in the deepest pools3-4 and 556 µS.cm in pool2 (variation 197/35%) (Fig. 4C), but the variations did not follow the topography of the cave. The same trend (min 454 µS.cm, max 553 µS.cm; variation 99/18%) was observed on the 2025 Piedras transect, but there the deepest pool10 showed the highest conductivity (Fig4-figure supplement 1).

Finally, we used PCA to evaluate the possibility of cave and pool signatures in an unsupervised manner (Fig. 4EF). Besides water parameters, here we included two additional geomorphological variables, pool depth and distance from cave entrance, for a more comprehensive description of each water body. The six caves studied clustered in the PCA in an almost non-overlapping manner (Fig. 4E), showing that each *Astyanax* cave represents a unique environment with a specific signature (ANOVA on pca1, F = 48.920, P < 2.2e-16; ANOVA on pca2, F = 7.899, P = 7.04e-6; twelve Tukey contrast tests out of the 15 possible were significant on pca1, while 5 out of 15 were significant on pca2). Moreover, clustering by pools (14 pools from 6 caves) revealed that while retaining the signature of their cave of origin, the different pools within a single cave have confidence ellipses that do not overlap (Fig. 4F), reinforcing the idea that each of the 14 pools studied constitutes a specific environment (ANOVA on pca1, F = 25.543, P < 2.2e-16; ANOVA on pca2, F = 68.433, P < 2.2e-16; forty-two Tukey contrast tests on the 91 possible were significant on pca1, while 56 on 91 were so on pca2). Together, these data suggest that the water conditions experienced by *Astyanax mexicanus* cavefish in different caves and in different pools within a single cave can vary significantly. In sum, the cave environment is not homogeneous, and each pool may be considered as a microhabitat.

### Hypoxia, a shared and extreme feature in cave waters

Cavefish have evolved hypoxia resistance through changes in the size or numbers of their erythrocytes, in the hypoxia inducible pathway gene regulation, and in gill morphology and the number of oxygen sensors (Boggs et al., 2022; Boggs and Gross, 2024; Boggs and Gross, 2025; van der Weele and Jeffery, 2022). To better understand the extent of hypoxia levels they face in their natural habitat, we measured oxygen content in the water of several caves in February 2022 and February 2023 (Fig. 5). As a control, we probed Río Choy (river location #2 on Fig. 1A), a well oxygenated running water located after a spring resurgence, where a large colony of surface *Astyanax* inhabit. We found 88.1% O2 saturation, corresponding to a dissolved O2 concentration of 7.02mg/L, which is consistent for a running river and acceptable for fish spawning and growth (Fig. 5B; light green). Of note, some still surface water bodies in the area may be less oxygenated. Conversely, in Pachón, Sabinos, Piedras, Chica and Subterráneo caves, we recorded strongly hypoxic conditions (Fig. 5A; orange to red).

**Figure 5:**
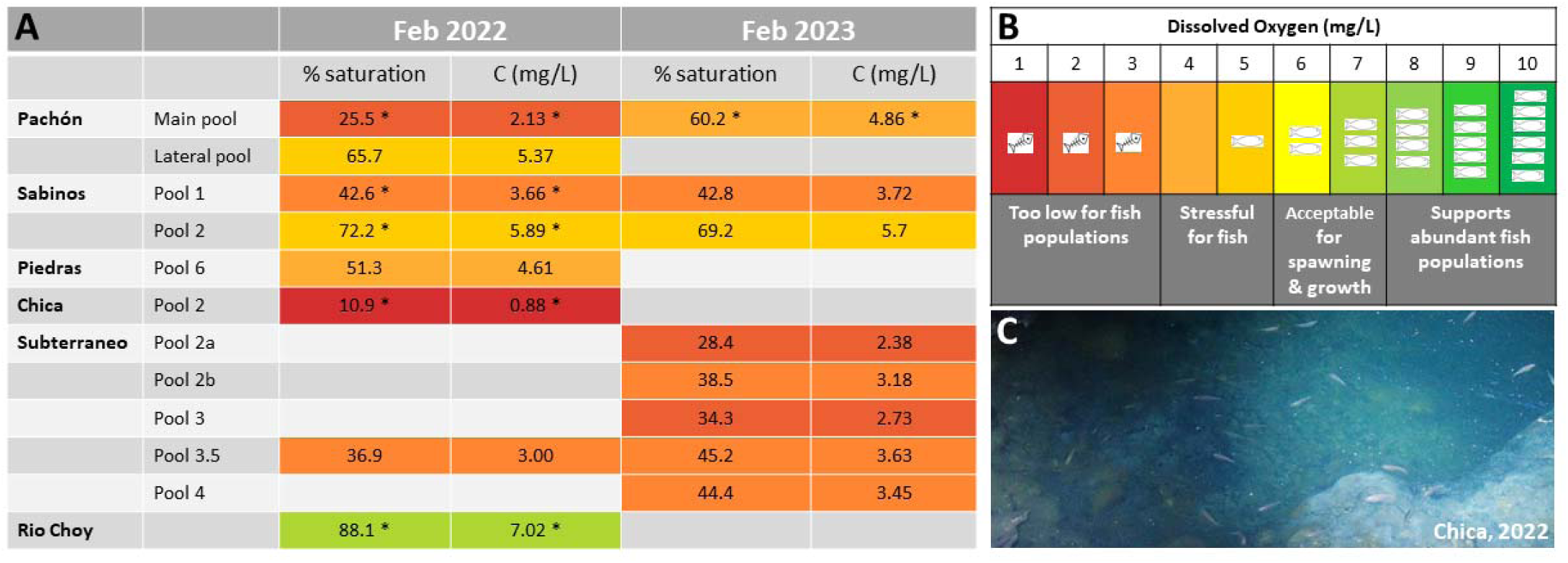
Oxygen content in caves and rivers. A: Data are from single measurements, except those with an asterisk (*), which are the mean value of 3 measurements made at 5 minutes intervals. B: Key color code for toxicity, adapted from Wikipedia. C: Picture of healthy-looking cavefish swimming in Chica, a strongly hypoxic cave.

We observed the most extreme conditions in Chica pool2 with 10% O2 saturation, i.e., 0.88 mg/L dissolved O2, which is in theory too low for fish populations to survive (Fig. 5B). A simple explanation for this dramatic situation is the presence of a very large bat colony in this cave, resulting in high levels of decomposing organic matter in the water (bat guano and cadavers, insects, leeches, ostracods…). Yet, Chica is inhabited by a large population of cavefish, which we have previously shown to be well fed, well grown, in good physical condition, and actually the biggest fish we have measured among all caves tested (Blin et al., 2020)(Fig. 5C).

All other cave pools tested showed low to very low O2 water content as well, all corresponding to lethal or very stressful conditions for “normal” fish (Fig. 5A-C). Interestingly, in Sabinos where we have recorded O2 two consecutive years (2022 and 2023), hypoxia seemed more pronounced in superficial pool1 (mean 42.7% O2 saturation; bat colony guano falling directly in the pool) than in deeper pool2 (mean 70.7% O2 saturation; protected from guano drops by low cave roof)(Fig. 5A), suggesting another level of heterogeneity between the two pools of this single cave. In this regard, it is noteworthy that the cavefish in Sabinos pool2 are significantly larger than those in pool1 (Blin et al., 2020)(Fig. 5C). Moreover, the presence of a fresh-water white sponge (Porifera) exclusively in Sabinos pool2 but not pool1 (Legendre et al., 2023a) could be explained by oxygen content as well, in addition to substrate and temperature differences.

Finally, an interesting observation was made in Pachón. The water of the main pool was strongly anoxic in 2022 (2.13mg/L dissolved O2) but less so in 2023 (4.86mg/L dissolved O2). This difference could be explained by the extremely low water level in Pachón main pool in February 2022, the lowest we have ever witnessed in this cave, with the fish swimming in a very reduced body of still water totally disconnected from the lateral pool (see (Legendre et al., 2023b) for pictures and see also below). This further suggests that cavefish not only have to cope with severe hypoxia, but also with fluctuating hypoxia levels over time.

### Annual longitudinal measurements of water parameters in caves

Having documented the global differences in physicochemical parameters of the water between surface streams and caves, between caves, and between pools within a cave, we next sought to investigate potential variations over the course of the year and the seasons. To this end, we installed and secured different types of probes (see Methods and Fig. 1B) in five selected caves: Pachón, Sabinos, Piedras, Chica (all Sierra de El Abra caves) and Subterráneo (Micos area). Probes installed in March 2019 were recovered in February 2022, after 3 years, due to Covid travel interruption. Those installed in February 2022-2023-2024 were recovered the next year, in February 2023-2024-2025, respectively. Some - but not all - of the probes provided full year records. In addition, to help interpreting the data from these probes, we set up camera traps near some cave entrances (Chica and Pachón) to better understand the hydrological regime and water flow into and out of the caves.

### Cueva de Los Sabinos

In Sabinos, we obtained records in pool1 and pool2, in 2019, 2022-23 and 2024-25 for temperature, conductivity and pH, including some in duplicates (Fig. 6 and Fig6-figure supplement 1). The average yearly temperature was 22.9°C in pool1 and 24.3°C in pool2 in 2019 (over 6 months), 24.0°C in pool1 and 24.3 in pool2 in 2022-23 (over 11-12 months) and 24.4°C in pool1 in 2024-2025 (over 12 months). The winter/spring values are very close to the one-off measurements shown above (22.5°C in pool1 and 24.4°C in pool2, n= 16 and n=18 point measures in February and March, respectively), and they confirm the temperature gradient in the cave, except during the rainy summer season (see below). Moreover, for both available recorded years, the temperature in pool2 was very stable (Fig6-figure supplement 1, turquoise curve/2019: min 24.24°C; max 24.36°C; var. 0.13; light green curve/2022: min 24.19°C; max 24.49°C; var. 0.30). Conversely, mild or more important and sudden temperature fluctuations were observed in pool1, both in 2019 (Fig6-figure supplement 1, dark blue curve, min 22.35°C; max 23.29°C; var. 0.94), 2022-2023 (Fig. 6A, blue, min 22.7°C; max 25.6°C; var. 2.9) and 2024-2025 (Fig. 6B, blue, min 22.8°C; max 26°C; var. 3.2). As observed from the 2022 recordings, pH (decreasing) and temperature (increasing) varied simultaneously but inversely in pool1 between February and June (Fig6-figure supplement 1). Both in 2022-23 and in 2024-25, conductivity varied inversely and synchronously with temperature (red curves; in 2022-23: min 459 µS.cm; max 637 µS.cm; var. 178; in 2024-25: min 390 µS.cm; max 599 µS.cm; var. 209) (Fig. 6AB). We interpret these data as pool1 being more exposed to the direct influence of rainy days and flooding than pool2, and having a more variable hydrological regime. In fact, in Sabinos pool1, in the 3 years recorded, temperature increased slowly and gradually starting in March-April (also seen on the 2019 partial curve). This was followed by sudden peaks of temperature increase occurring during the summer (16 August and 6, 20, 27 September 2022; 22 June, 1, 11, 19, 26 July and 15 September 2024), in a reproducible pattern in 2022-23 and in 2024-25 rainy seasons. Synchronously, sudden drops in conductivity occurred (red curves), indicating rainwater with low mineral content entering the cave. Thus, due to influx of surface water during the 3 months of summer, temperature of pool1 became higher than in pool2, transiently inverting the temperature gradient we have previously described with point measurements obtained in February/March (Fig. 4A).

**Figure 6:**
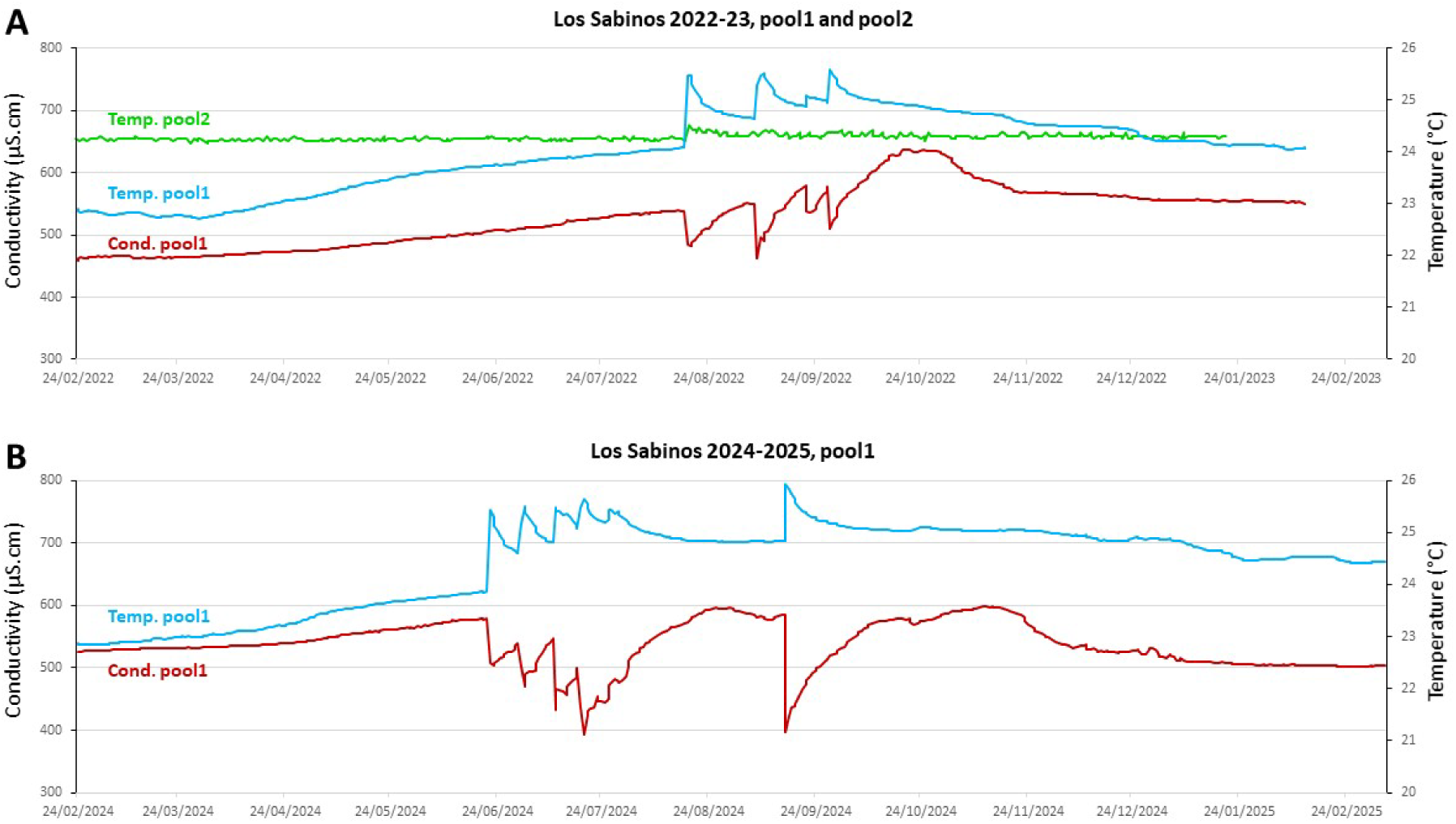
longitudinal measurements of water parameters in Sabinos. A: Evolution of temperature and conductivity in Sabinos pool1 and pool2 from February 2022 to February 2023. B: Evolution of temperature and conductivity in Sabinos pool1 from February 2024 to February 2025. Color codes for recorded parameter, pool and year of interest are indicated. Note that Pool2 is globally warmer than Pool1 except during the rainy season; that Pool1 seems to undergo more fluctuations than Pool2; and that temperature and conductivity parameters vary in an inversely correlated manner in Pool1 at a daily timescale.

Importantly, on 16 August 2022, when a massive and sudden influx of water flooded the cave as deduced from the 1.39°C temperature increase observed on a single day in pool1 (blue curve), a smaller but significant and synchronous peak (+0.2°C) occurred the same day in pool2 (green curve, Fig. 6A). These data strongly suggest that the flood of that day penetrated to the bottom of the cave, and that pool1 and pool2 became connected at least during a few days. These observations have important outcomes for Sabinos cavefish biology (and other inhabitants of the cave, arthropods, bats and sponges (Elliott, 2018; Legendre et al., 2023a)). First, they are sporadically exposed to catastrophic and stressful events of environmental change, to which they must react and adapt. Second, the cavefish populations of pool1 and pool2 can mix from time to time, which means that genetic differentiation cannot occur between pools and the Sabinos cavefish must be genetically homogeneous.

### Cueva del Río Subterráneo and Cueva Chica

Subterráneo and Chica are two caves in which introgression of surface fish swimming in surface streams occurs regularly, thus they host hybrid fish populations (Bibliowicz et al., 2013; Breder, 1942).

In Subterráneo, surface fish are washed into the cave entrance in the rainy season, as indicated by their abundance in the first “perched” pool of this cave, and their presence further down in the cave where they co-habit and breed with cavefish. Other epigean species of Cichlidae and Poeciliidae as well as crayfish are regularly observed. We have personally witnessed the violent flooding of this cave, so seasonal variations in water parameters were expected. In pool2, we obtained one year of records of temperature (in triplicate), pH and conductivity (Feb2022-Feb2023) (Fig. 7A). The temperature curves recorded in triplicate with three different types of instruments (different blue shades on Fig. 7A) showed identical profiles, suggesting that the data we present in this paper are consistent and reliable. As anticipated, throughout the year, Subterráneo pool2 was subjected to strong, sudden and synchronous variations in temperature (min 19.8°C; max 28.2°C; var. 8.4), pH (pink curve, min 7.15; max 8.64; var. 1.49) and conductivity (red curve). Note that we consider the drop of conductivity to almost zero in August 2022 to be an artifact of uncertain origin (the probe probably got transiently out of the water). After a gradual increase in temperature between February and August 2022, several sudden peaks in temperature and drops in conductivity occurred on 17 August, 5 and 8 September and later, followed by progressive return to more stable winter values in December (Fig. 7A). The increases in water temperature in August and September are consistent with water flowing into the cave from the surface after heavy and warm summer rains. We interpret the December drop to 20.88°C, associated to a moderate conductivity drop, as cold winter rainwater entering the cave (Fig. 7A). Air circulation may contribute to the regulation of temperature after these fluctuations and during the rest of the year. Overall, the massive variations in water parameters observed in Subterráneo suggest a drastic, complex and dynamic hydrological regime, to which the resident cavefish are exposed.

**Figure 7:**
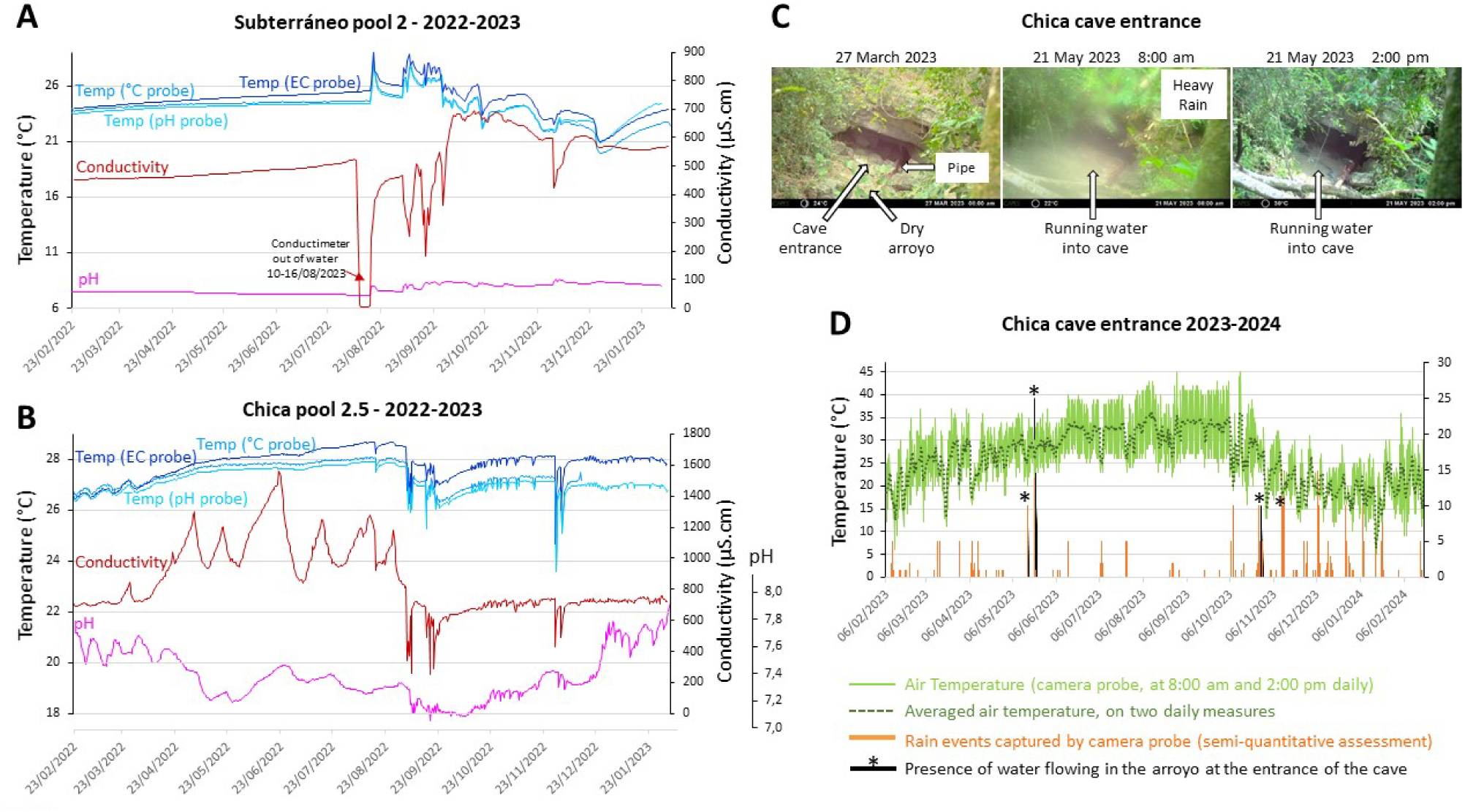
longitudinal measurements of water parameters during one year in Subterráneo and Chica. A: Evolution of temperature, pH and conductivity from February 2022 to February 2023 in the Subterráneo cave pool2. Color codes for recorded parameter are indicated. Note that water temperature recorded from the specific temperature probe, or the conductivity probe (EC), or the pH probe show identical results and temperature variation patterns. We interpret the sudden, unexplained and unrelated drop in conductivity between the 10 and the 16 October 2023 as the device being out of water. After this event, conductivity records resumed and are consistent as they are correlated to variations in temperature and pH at a daily timescale. B: Evolution of temperature, pH and conductivity from February 2022 to February 2023 in Chica pool2.5. Color codes for recorded parameter are indicated. C: Examples of pictures captured by the camera trap facing the entrance of the Chica cave between February 2023 and February 2024. The dates are indicated. D: Evolution of air temperature and water flow regime and rain at the entrance of the Chica cave, as captured by the camera device installed on a tree facing the cave entrance, between February 2023 and February 2024.

In Chica, it is believed that surface fish from the nearby Río Tampaón could enter the cave through the deepest part of the cave, rather than through the entrance used by humans at ground level. In this cave, in pool2.5 (a medium size pool between pool2 and pool3) we obtained one year of records of temperature (in triplicate, identical curves in blue shades), pH (pink) and conductivity (red) (Feb 2022-Feb2023) (Fig. 7B). The longitudinal measurements confirm that Chica is the warmest cave in the area (average 27.4°C). They also show that the annual variations are important for the three parameters recorded, especially for conductivity and temperature (temperature: min 23.6°C, max 28.1°C, var. 4.5; pH: min 7.07, max 7.95, var. 0.88; conductivity: min 252 µS.cm, max 1560 µS.cm, var. 1307). In Chica like in other caves described above, the temperature increased progressively from February to August (Fig. 7B). However, contrarily to other caves, this gradual increase of temperature was associated to large oscillations in water conductivity (increases, not drops) and pH, suggesting that these variations may not be related to water entrance. We hypothesize that they could be the result of chemical and organic interactions in the water, mainly driven by the activity of bats, which are present in huge numbers in this cave, and which contribute to eutrophication of the milieu. Then, from the end of August, the temperature oscillated and underwent several sudden, transient and significant decreases (17 Aug, 5/7/16/20 Sept, 1/5 Dec), fully synchronized with transient decreases in conductivity indicating rainwater entrance. Such events of Chica cave flooding from the dry “arroyo” at its entrance was not clear from discussions with the local farmers. Therefore, we set up a time-lapse camera focused on the cave entrance and obtained a year’s record of rainfall and outside air temperature, from February 2023 to February 2024 (Fig. 7C). Rainy days and the resulting water regime were easy to identify and interpret (Fig. 7CD). Heavy rains caused the arroyo to flow actively and rainwater to enter the Chica cave on a massive scale four times in 2023 (twice in May, twice in October-November), confirming the hypothesis that rainwater does enter in Chica by its main entrance. Our direct observation of several *Astyanax* surface fish in the small pool0 close to the cave entrance in March 2017 and February 2023 is also consistent with this finding. Taken together, our data suggest that Chica cavefish are exposed to the most extreme and changing hydrological regime of any *Astyanax* cave we have studied.

### Cueva de El Pachón

Pachón is the most studied *Astyanax* cave, for both laboratory and field research (Blin et al., 2020; Espinasa et al., 2017; Espinasa et al., 2023; Hyacinthe et al., 2019; Krishnan et al., 2020; Legendre et al., 2023b; Ornelas-Garcia et al., 2018; Peuss et al., 2020; Simon et al., 2017), so understanding its ecology is paramount (Fig. 8).

**Figure 8:**
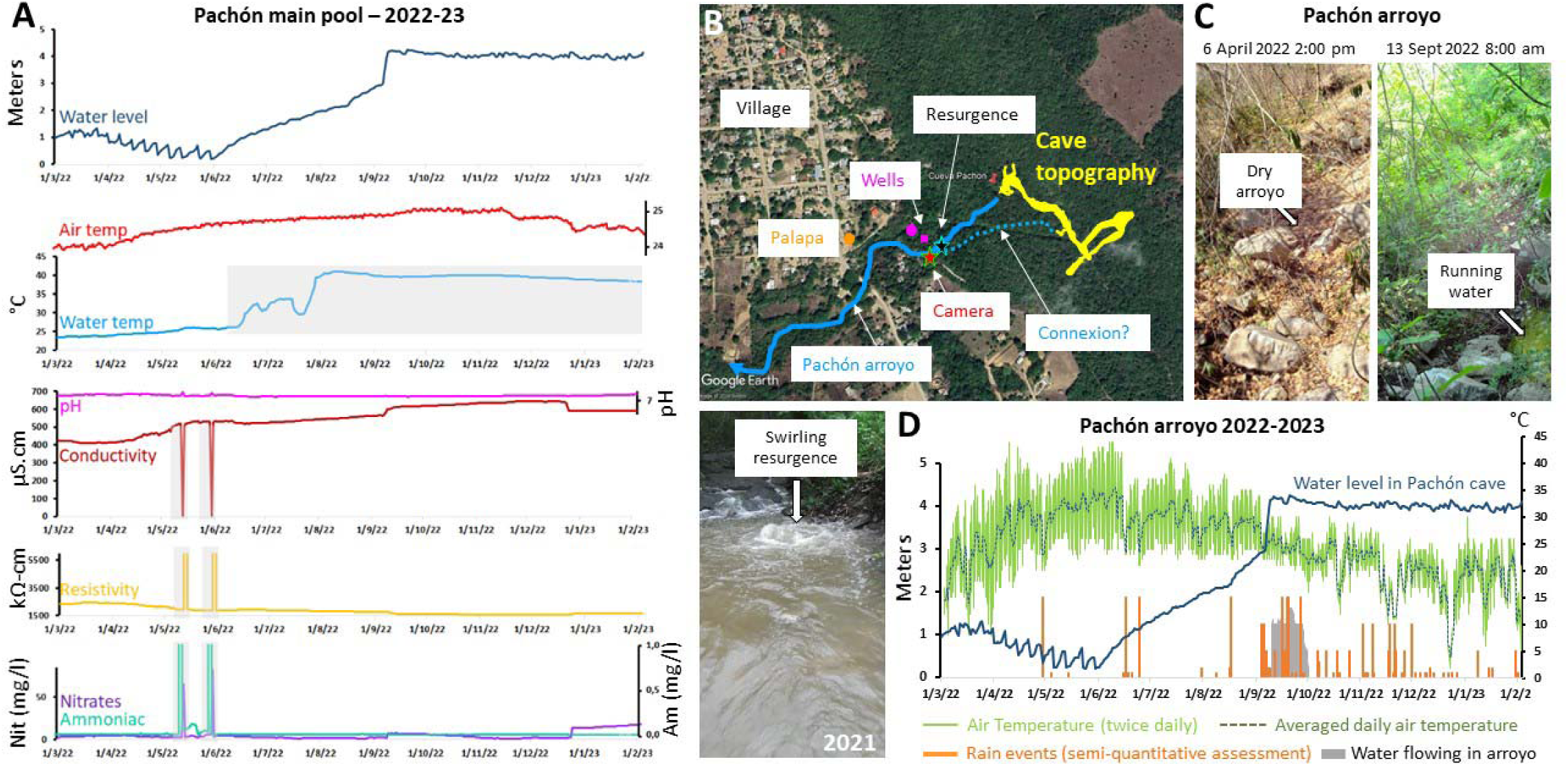
longitudinal measurements of water parameters during one year in Pachón. A: Evolution of physico-chemical water and air parameters from February 2022 to February 2023 inside the Pachón cave, main pool. The grey-shaded parts of the curves are interpreted as artifacts and should not be considered, because they result from the probes being out of the water, when levels reached extremely low values (see text). Color codes for recorded parameter are indicated. B: Schema of the geography around the Pachón cave, superimposed on Google Earth view of the area. Relevant information is labelled. The bottom photograph taken by a local in 2021 shows the active resurgence, with important water flow. C: Examples of pictures captured by the camera trap installed along the arroyo flowing below the Pachón cave entrance. The dates are indicated. The position of the camera is shown in panel B. D: Evolution of air temperature, rain and water flow regime at the level of the arroyo below the Pachón cave entrance, as recorded by the camera device installed on a tree facing the arroyo. The water level recorded by multi-parametric probe inside the cave (panel A) is superimposed in order to show the correlation in time between the different events.

In 2019, we recorded temperature in the Pachón main and lateral pools. In sharp contrast to other caves, water temperature was very stable (main pool/4months, min 24.4°C, max 25°C, var. 0.6; lateral pool/9months, min 24°C, max 25.4°C, var. 1.4) (Fig8-figure supplement 1) and showed only a minor and continuous increase between March and September. The first months of the 2022 records confirmed this trend (Fig. 8A, blue curve). We propose that these minor variations in water temperature follow the evolution of the cave air temperature (red curve on Fig. 8A, recorded in 2022-2023: min 23.9°C, max 25.1°C, var. 1.2), which in turn is influenced, albeit strongly buffered, by seasonal variations in outside air temperature (Fig. 8D, green curve). This scenario is consistent with Pachón being a perched cave that is not flooded by rivers that overflow during the rainy season, and that receives water mainly by percolation through the epikarst.

In February 2022, when water was extremely low in the cave as mentioned above, we setup a multi-parametric probe at the back of the main pool (map in Supplemental File1). Water level was included in the recorded parameters (dark blue curve on Fig. 8A). To our surprise, between March and May 2022, the water level continued to fall by another meter, with very regular weekly oscillations (n=12) between the 9^th^ March and the 30^th^ May, every seven days on average, which we are tempted to interpret as pumping episodes through the pipe installed there. For the cavefish population swimming there, the amount of water available must have been minimal and the situation extremely dangerous for at least two months. After this “catastrophic” event, the water level rose by more than 4 meters in about 4 months, returning to the "normal" water level that we and others have repeatedly observed when visiting the cave at different seasons (including when we recovered the multi-parametric probe in February 2023). This shows that Pachón main pool occasionally undergoes variations of hundreds to thousands of cubic meters of water volume. Before February 2022, the last known record of such low water in Pachón was in March 2003, when Jeffery, Yamamoto and Espinasa took this opportunity to discover and survey the Maryland extension galleries.

Unfortunately, one consequence of the extremely low water levels is that our temperature records cannot be considered reliable after ∼mid-May 2022 (blue curve, shaded in Fig. 8A), because the disposable sensors of the multi-parametric probe is likely to have been partially dried/damaged. Likewise, the two peaks observed simultaneously on the conductivity, resistivity, ammoniac and nitrate curves are likely artefacts due to very low water levels. However, the available records of pH, conductivity and resistivity (which mirror each other, therefore considered reliable), as well as nitrate and ammonia, suggest that water conditions in Pachón were relatively stable throughout the year, despite the large amount of water that replenished the water chamber.

Finally, locals caught our attention on strong episodes of water resurgences at the bottom of the dry arroyo under the Pachón cave (named “El Venero” - the spring - by locals; Fig. 8B). This suggested that when water level is high, the cave might discharge into the arroyo *via* this resurgence (at least partially). We installed a time-lapse camera to capture the rainfall and the water regime in this arroyo from February 2022 to February 2023 (Fig. 8CD). The data collected show numerous rainy days in September 2022 (orange on Fig. 8D), associated with a period of about a month during which water flowed in the arroyo (grey on Fig. 8D), fed by the resurgence (Fig. 8CD). Strikingly, this period coincided with the time when there was an abrupt raise of about 1.5 meter in water level in the Pachón cave pool (Fig. 8D, dark blue curve superimposed). We interpret these data as the heavy rain contributing to cave pool recharge through percolation, followed by cave pool overflow into karst cavities and into the network connected to the resurgence at the level of the arroyo (Fig. 8B). The outflow siphon at the end of the lateral pool (also known as pool1 or corridor pool) would be directly or indirectly connected to El Venero resurgence. It follows that the water level in Pachón cannot be higher than a certain threshold (about 4 meters above the level of our multi-parametric probe) or it will overflow. It also means that some cavefish may be washed out of the cave during these episodes, further contributing to the decline of the population in this cave (Legendre et al., 2023b).

### Sótano de Las Piedras

Among the caves we have studied yearlong, Piedras is the only one with a pit entrance, about 20 meters deep (Elliott, 2018). As mentioned above, this topography probably explains why the first pools of this cave display cool water temperatures, as measured in February-March (Fig. 4). In addition, we obtained two complete years of recordings, in 2022-23 in pool5, and in 2024-2025 in three different pools (Fig. 9). Both years, winter temperatures in pool5 (light blue curves) were consistent with previous point measurements, i.e. between 19-20°C at the lowest, and showing rather cool temperatures throughout seasons. Moreover, the 2024-2025 recordings showed that despite fluctuations during the rainy season, the increasing temperature gradient from superficial to deeper pools was maintained throughout the year (mean year temperatures: pool5 22.1°C, pool9 23.2°C, pool10 24.4°C), confirming the tendency observed from point and transect measurements in February-March (Fig. 4A,D).

**Figure 9:**
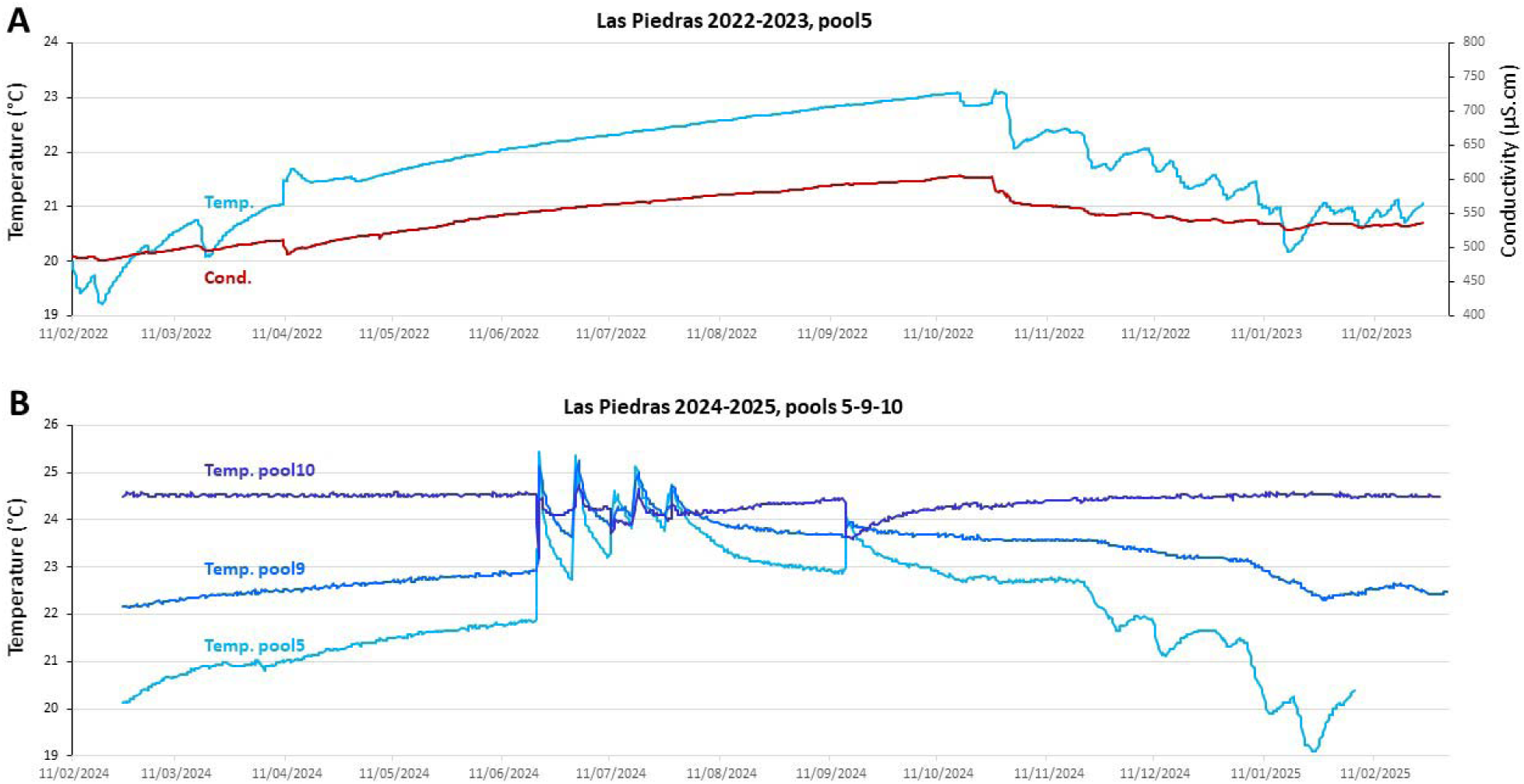
longitudinal measurements of water parameters in Piedras. A: Evolution of temperature and conductivity in Piedras pool5 from February 2022 to February 2023. B: Evolution of temperature in 3 different pools of Piedras from February 2024 to February 2025. Color codes for recorded parameter and pools are indicated.

During 2022-23, temperature and conductivity in pool5 showed modest and regular fluctuations without sudden changes during the rainy season (temperature: mean 21.8°C, min 19.9°C, max 23.1°C, var. 3.2; conductivity: mean 548 µS.cm, min 486 µS.cm, max 606 µS.cm, var. 119), suggesting stability of conditions inside the cave that year (Fig. 9A). In 2024-25 however, six sudden changes in temperature were observed, synchronously in pools 5-9-10 (21 June, 2-13-19-28 July, 15 September 2024) (Fig. 9B). On the 21 June for example, temperature in pool5 increased by 3.56°C (from 21.87 to 25.43°C; light blue curve), temperature in pool9 increased by 2.23°C (from 22.9 to 25.13°C; blue), and temperature in pool 10 decreased by 1.33°C (from 24.53 to 23.2°C; dark blue) (Fig. 9B). We interpret these variations as warm summer rainwater flooding the cave, leading to an increase in pool5 water temperature and water level, and leading to an overflow of pool5 into the next pools, and so on. When flooding reaches pool9, it also induces an increase in temperature because pool9 is initially cooler than rainwater. Conversely, when flooding reaches pool10, the “hottest” pool of Piedras, it causes a temperature drop. The six heavy rain episodes that occurred in 2024-2025 induced the exact same pattern of temperature changes in the different pools, suggesting the possibility of important water mixing as well as cavefish mixing between pools, at least downwards. Indeed, the cave topography and conformation with limestone “dams” and a “slide” ahead of pool9 seem incompatible with water and fish going upwards; and pools 1-5 are most probably stable over years, as deduced from well-formed rimmed concretions on rocks at water level. Interestingly, after each rain episode, in each pool temperature rapidly tended to return to values closer to average specific temperature values (i.e., re-decreased in pool5 and pool9, and re-increased in pool10), suggesting that the local configuration around each pool has a strong influence on temperature regulation. Important factors may include fresh air circulation close to cave entrance, localization of bat colonies in the deepest part of the cave, and thermal exchanges with the rock substrate, which must have an important thermal inertia.

Finally, in both recorded years, pool5 (but not pools 9 and 10) showed slower and smaller temperature fluctuations from November to February (Fig. 9AB), accompanied by a modest decrease in conductivity (Fig. 9A). These observations and the similarity between the 2022-23 and 2023-24 curves suggest that the anterior-most part of the cave is under aerologic influence during the autumn and winter “cold” period, due to the special conformation of the deep pit entrance, with air currents contributing to the cooling of the water temperature. Alternatively, we cannot rule out that occasional autumn and winter “cold” rain reaches only the first pools of the cave, and contribute to the negative regulation of their temperature.

In sum, our dataset suggests that the hydrological regime in Piedras is variable from a year to another. The relatively low numbers of cavefish we have repeatedly observed in the clear water of pool5 and the larger numbers of cavefish we have observed in pools 9-10 with abundant organic matter accumulation may result from the episodic washing of superficial pools into the deeper ones, in a sort of cascade, or domino effect.

## Discussion

The ecology, hydrodynamics and classification of groundwater ecosystems is a subject of research and attention (Aquilina et al., 2023; Robertson et al., 2023), especially in the interest of groundwater ecosystem functions and services (Hose et al., 2023; Kretschmer et al., 2023). This report is, to our knowledge, one of the few to document and interpret cave water quality and hydrodynamics for an integrated view of cave biology and the evolution and conservation of cave organisms (Koren et al., 2023), and the first for *Astyanax* (Malard et al., 2023; Torres-Paz et al., 2018). The point and longitudinal measurements of water parameters that we obtained in several caves and rivers hosting populations of *Astyanax mexicanus* between 2009 and 2025 have profound implications for understanding the evolution of cavefishes and the biology of the species in general.

### Macrohabitat

The surface fish *Astyanax mexicanus* swim in subtropical freshwaters in Mexico and as far north as Texas. Conversely, the cave-adapted morphs of the species inhabit a series of 33 caves in a restricted region of North-East Mexico (Elliott, 2018; Espinasa et al., 2018; Espinasa et al., 2020; Mitchell et al., 1977). Water quality sampling in this area, in surface rivers, ponds, springs on the one hand and in caves on the other, shows marked differences between epigean and hypogean waters and reveals strong signatures of these two macrohabitats: cave water is fresher, much less conductive than surface water, and highly anoxic. Our large-scale dataset corroborates earlier punctual and sparse reports on these variables (Breder, 1942; Elipot et al., 2014b; Ornelas-Garcia et al., 2018; Tabin et al., 2018).

Darkness and the absence of primary production are generally considered the hallmarks of the cave environment. However, water quality may also play a very important role as an environmental variable driving the evolution and adaptation of cavefish, and as a *bona fide* contributor to this so-called “extreme” habitat. In fact, the large shift in water conductivity from surface to cave has been proposed as a potential abiotic stress factor that could have triggered an HSP90-related stress response and unmasked cryptic variation, allowing the initial evolution of eye loss during cave colonization (Jarosz and Lindquist, 2010; Rohner et al., 2013). To our knowledge, low levels of dissolved oxygen have not been tested in this direction, but it is hard to imagine that the severe and sudden hypoxia encountered by the first cave dwellers did not have any effect on their stress response and adaptation.

*Astyanax mexicanus* belongs to the order Characiformes and the family Characidae. Yet, before evolving the specific hypoxia-resistant traits that extant cavefish possess today (Boggs et al., 2022; Boggs and Gross, 2024; Boggs and Gross, 2025; van der Weele and Jeffery, 2022), they must have survived in conditions that are lethal to highly stressful for most fish. This may indicate that *Astyanax mexicanus*, as a species, has hidden characteristics that enable it to withstand these harsh conditions. If true, such a preadaptation to hypoxia, together with other traits like the ability to breed in the dark (Simon et al., 2019), would also explain why only *Astyanax* has colonized and thrived in Mexican caves. In fact, fish from other families like Poeciliidae and Cichlidae that we occasionally observed in these same caves (regularly at least in Chica and Subterráneo) are always in poor condition, they do not survive and do not form established populations in the *Astyanax* caves network (personal observations). An alternative possibility is that, when surface fish colonized caves around 20.000 years ago (Fumey et al., 2018; Policarpo et al., 2021), cave water was slightly more oxygenated and they did not have to cope with anoxia on top of darkness. Notably, we also witnessed surface fish swimming in water with a pH of 9.7 and a temperature exceeding 30°C, suggesting that the extant species (and ancestor) as a whole is quite resilient and plastic to environmental water conditions.

### Mesohabitat

Every cave is special and unique. Besides the low conductivity of the subterranean waters that is a shared feature of all caves studied, other parameters can vary significantly between locations. One of the most important may be the temperature of the water. The difference between Chica, the hottest cave at ∼27-28°C, and Arroyo, the coolest cave at ∼16-17°C, is large and must have important consequences for reproduction and development, metabolism, energy homeostasis and behavior of the cavefish populations – as they are poikilothermic, i.e. their body temperature varies with the ambient temperature eg (Zahangir et al., 2022). The origin of such differences probably lies in the general geological and biological characteristics of the caves. In Chica, for example, the tens of thousands of bats that live there act as a powerful air heater, while the entrance to the very large and long Arroyo Cave is located at the bottom of a 25-meter pit embedded in the coolness of the forest. Many other combinations are possible, and air dynamics linked to cave geomorphology may have a certain influence. Of note, the capacity of cavefish to survive at variable temperatures is probably inherited from surface ancestors, as we have found that extant surface fish can experience temperature ranges between 19°C and 31°C.

Interestingly, river-dwelling *Astyanax* and cavefish from the Tinaja and Molino caves have a temperature preference that is significantly higher (28.3°C) than Pachón cavefish (25°C), as measured in behavioral assays (Tabin et al., 2018). The difference between surface and cave fits well with the fact that temperatures in rivers are higher than in caves, even at the end of winter when all our point measurements were obtained. The difference between caves is also consistent with our dataset and suggests that local adaptations may have occurred in different cave populations in response to specific environmental water cues. It would be interesting to address the genetic determinism of the physiological and behavioral adaptations present in, e.g., the Chica and the Arroyo populations discussed above (Tabin et al., 2018). In the same vein, it would be important to study the genetic basis of hypoxia resistance, as the physiological changes needed to survive in water with 10% O2 saturation (Chica) are likely to be different from those needed to survive in water with more than 60% O2 saturation or fluctuating O2 levels (Pachón, Sabinos) (Boggs et al., 2022; Boggs and Gross, 2024; Boggs and Gross, 2025). Moreover, water parameters and particularly oxygen levels - rather than the cave vs surface environment - seem to influence *Astyanax* gut microbiome, which has essential roles in the regulation of metabolism, physiology and health (Ornelas-Garcia et al., 2018).

Another aspect that makes each cave different is the variety of seasonal and multi-annual hydrological regimes. From our longitudinal multiparametric measurements, it appears that water quality can be either stable (Pachón) or vary slightly or more significantly throughout the year, either gradually or suddenly (other caves). Some caves receive water mainly from outside precipitation (Sabinos, Subterráneo), or from percolation (Pachón), or from both precipitation and nearby streams (Chica). Some caves are perched and drain into the hydrological system, so their water level has a maximum (Pachón), while others can be completely flooded. In addition, in the caves where a pipe is installed such as Pachón and Chica, the water level and the amount of water available may in principle be subject to very strong fluctuations due to anthropic activities, which we did witness in both cases (very low levels in Pachón in 2022 and Chica in 2024). Thus, qualitatively and quantitatively, the hydrodynamics of the studied caves are different - although globally the discharge and the recharge are very fast, because karstic systems can be flashy by nature (Aquilina et al., 2023).

Incidentally, caves in the Micos area (e.g. Subterráneo) that are flooded after rainfall are likely to receive large amounts of phytosanitary chemicals from nearby sugar cane plantations, a situation that is detrimental to cave fish and other cave animals, as well as to the quality of the underground water. Our qualitative observations that fish from Subterráneo cave are more infested by parasites than in other caves are in line with the recent reports by Santacruz et al. (2023). In this study, Subterráneo cavefish were the most infested by parasites among the 18 cave populations tested, with the presence of 9 parasites taxa in the cave and 3 parasites per individual fish on average, and the unique presence of the invasive anchor worm *Lernaea cyprinacea* (Santacruz et al., 2023). In principle, caves that are not flooded escape this threat (e.g. Pachón, only 2 parasites taxa (Santacruz et al., 2023)). To our knowledge, the impact of chemical pollution of water has not been studied in *Astyanax* caves, but we expect it to be an important conservation issue (Boulton et al., 2023; Hose et al., 2023).

The complex and specific hydrodynamics we have uncovered make the cave environment less stable and homogeneous than commonly thought, at least from the perspective of cavefish. Researchers who enter caves for fieldwork usually do so at the end of the dry season for obvious safety reasons, so they have a biased view of what cave life is. In fact, cavefish experience daily (driven by the entry and exit of bats at dusk and dawn (Beale et al., 2013)), seasonal (driven largely by rainfall and outside air temperature), climatic (hurricanes hit the area sporadically throughout decades or centuries) and other non-natural regular or irregular rhythms (driven by human activity such as pumping of water, mining activities…). Different caves are differently impacted by these four types of influences, depending on their configuration and geology (Elliott, 2016; Elliott, 2018). Hence, each cave environment is unique, and each cave population has evolved according to different selection pressures (besides the critical and shared absence of light). Although the causal relationships between specific environments and local cavefish genetics will probably always remain elusive, the different conditions found in different caves may have contributed to shaping slightly different evolutionary trajectories (Borowsky, 2008; Policarpo et al., 2024).

Another consequence of our findings is the need for cavefish to cope with stress associated to fluctuating environment (Cobham and Rohner, 2024). Some authors have described reduced behavioral stress responses in cavefish populations compared to surface fish with a novel tank test (Chin et al., 2018), and they hypothesized that this corresponds to the transition between predator-rich epigean and predator-free cave environments. However, according to the present data, we would argue that the environment of the cave is not free of stress. On the other hand, others have described a mutation in the *MAO* gene (Elipot et al., 2014a), which is distributed in all the cavefish populations of the El Abra region, and which confers low basal anxiety to fish in stable conditions but also strongly increases the stressability of fish in response to a novel tank (Pierre et al., 2020). These results are consistent with cave life being predator-free and rather stable most of the year, as well as with the need to react fast and efficiently to sudden and sometimes cataclysmic events that occur during the rainy season. As year-long records of cavefish behaviors are technically impossible to obtain in most caves, indirect monitoring of their activity and distribution in the caves through underwater long-term passive acoustic recordings would bring some valuable insights on how they cope and react to seasonal changes in each cave (Aalbers and Sepulveda, 2012; Lammers et al., 2008).

### Microhabitat

Finally, our data suggest that within a cave, each pool is different. Again, one important variable is temperature, which shows a low-to-high gradient from more superficial pools to pools further away from cave entrance in a reproducible manner in all caves, at the end of winter. Differences can be important, such as ∼5°C between pool5 and pool10 in Piedras, or ∼7.5°C between Subterráneo pools. As cavefish seem to have temperature preference that are different between caves and that this trait is genetically encoded (Tabin et al., 2018), our findings raise the question of how this temperature preference is established and regulated at cave population level. In Tinaja, different pools also have different conductivities and in Piedras, different pools show different pH.

In addition to water parameters, another variable is nutrient availability, and we have observed that different pools within a cave vary by biotic and abiotic aspects. For example, a part of Sabinos pool1 - but not pool2- is located under a bat roost, and the spatial distribution of cavefish in the small part of pool1 where bat guano droppings are available suggests that this parameter is very important (Espinasa et al., 2024). Conversely, Sabinos pool2 -but not pool1-contains freshwater sponges (Legendre et al., 2023a), so the chemical signature of the water including numerous sponge metabolites must be very specific. As another example, some pools have a muddy substrate (Tinaja pool2) while others have a rocky substrate (Tinaja pool1), thus offering different compositions in microorganisms.

Our analyses further show that whereas cave pools are separated when researchers visit them, they connect from time to time when water level raises. In Pachón, fish residents of main and lateral pool may be separated for several months, but they will eventually mix, at least once a year, when water level raises. In Sabinos too, where pool1 and pool2 are distant from about 150 meters and at distinct depth, mixing occurs after heavy rain and flooding. This is consistent with the lack of genetic differentiation between pools within caves (unpublished results). Moreover, such mixing may result in the redistribution of fish individuals and populations between pools from a year or a month to another, again generating stress, and forcing them to be plastic and re-explore their environment to map their novel swimming space - a task achieved without vision, as always.

### Conclusion

Cave colonization by surface *Astyanax* occurred about 20,000 years ago during glacial times, twice independently in the Micos area on the one hand and in the Sierra de El Abra/Sierra de Guatemala region on the other (Policarpo et al., 2024). At that time, the biotic environment, the geography, the geology/topography, the climate, the hydrodynamics and the water quality must have been different from today. In any case, the greatest challenge for the first cavefish was the absence of light. However, in the course of their progressive colonization of karstic cavities in a “stepping stone” mode (33 caves inhabited by *Astyanax* are known today) they have encountered different, specific, sometimes extreme conditions in water regimes and qualities that have undoubtedly shaped the evolutionary trajectories of the different populations.

## Methods

### Geographical area studied

The stations studied are located in the San Luis Potosi and Tamaulipas regions of northeastern Mexico (part of the Sierra Madre Oriental), an area approximately 130 km long and 60 km wide. Between 2009 and 2025 we collected data on water physico-chemical parameters from 13 caves and 30 surface stations (Fig. 1A). Some stations were surveyed many times, with different interval times (days, months, years). All point measurements are reported in Supplemental File2 (raw data file) together with some additional relevant information. Moreover, longitudinal recordings were performed in a subset of cave stations.

All cave stations were inhabited by *Astyanax* cavefish (or hybrids), and all surface stations were inhabited by *Astyanax* surface fish, except one, station #6 at the Hotel Taninul hot sulphur swimming pool. Values from station #6 were not included in graphs and analyses, but are worth reporting, as they are part of the local hydrological system. There, temperature was 36.7°C in 2009 and 37.8°C in 2011, pH was 6.8 in 2009 and 6.95 in 2011 and conductivity was 1276 µS.cm in 2011.

### Recording instruments

We used different methods and apparatuses to record water parameters (Fig. 1B).

#### Point measurements

One-off measurements were taken using a Combo Hanna HI98129 meter (Hanna Instruments, USA) for pH, electrical conductivity and temperature. The Combo was calibrated regularly in the laboratory and in the field using the HI98129 manufacturer’s protocol. The Combo sensor for pH was changed as often as necessary (Hanna Instruments, USA).

Some additional pH controls were performed with colored Tetratest (TETRA, Copyright © Spectrum Brands, Inc., USA, and JBL GmbH & Co., Germany). All colored test used were recently purchased and were open only for the fieldtrip.

A field oximeter Hanna HI98193 (Hanna Instruments, USA) wired with HI764073 probe was used to measure dissolved oxygen/temperature.

#### Longitudinal measurements

For long-term measurements, in most caves we used HOBO probes. Water or air temperature measurements of 2019, 2022, 2023 and 2024 were made by MX2201 and MX2202, pH/temperature by MX2501, and electrical conductivity/temperature by U24-001 (Onset HOBO® Data Loggers, USA).

A multi-parametric probe AQUATROLL 600 (type 74050, IN-SITU©2022 In-Situ Inc., USA) was used for longitudinal measurements in the Pachón cave (as well as for some point measure mainly in February 2022 and March 2025 in other caves). This probe is designed with four smart sensor ports for interchangeable sensors, an LCD screen and Bluetooth connectivity. For this study, it was fitted with four sensors as follows: one for pH/ORP (758768), one for Nitrate NO3-N (ISE, 742828), one for Ammonium NH4+N (ISE, 704780) and one for electrical conductivity/temperature (778296). Details on probes resolution, accuracy, range and calibration solutions are available in the manufacturer’s instructions and technical sheets.

To configure the probes and retrieve data, we used a mobile Bluetooth connection with mobile applications designed specifically for these probes. The VuSitu app (In-Situ, Inc., version 1.19.12) was used for AQUATROLL 600. The HOBOmobile app (ONSET, HOBOmobile® for Android, version 2.0 edition: 1029 ©2019 Onset Computer Corporation) was used for the HOBO MX2201, MX2202, and MX2501 probes. The HOBOware software (ONSET, HOBOware®, version 3.7.19, ©2002-2020 Onset Computer Corporation) was used for the U24-001 probe, which requires an optical connection through the Base-U-4 Optic USB Base Station (Onset HOBO® Data Loggers, USA) to restore or configure the probe. We programmed, configured, and verified the probes in the field just prior to deployment. The majority of probes have logging rates ranging from 1 second to 18 hours. We choose 12 hours for longitudinal studies.

Finally, to record time-lapse photographs of rainy/sunny days and outside air temperature, we used TimelapseCam Pro (Wingscape, WCT-00126).

#### Transect measurements in Piedras pools

On 3 March 2025, we used the multi-parametric AQUATROLL 600 probe to record temperature, pH and conductivity systematically in all the 10 successive pools of Piedras. Measures were taken twice in each pool, once while entering and going down the cave, and once on the way back exiting the cave. Values were identical or highly similar for the two replicate measurements, except for pH records in pools 8, 7, 6 and 5. There, pH values were slightly lower for the second measurement, when going up (0.15 to 0.2 pH variation). We interpret this minor variation by the mixing and swirling of water due to experimenters walking or swimming in the pools on their way down, causing CO_2_ degassing, O_2_ exchange, and organic matter resuspension that can affect pH but not temperature nor conductivity.

### Installation and fixation of probes

Fixing the longitudinal probes/data loggers was a challenge. The probes had to be small, light, and easy to attach without any prior information about the water flow and the potential debris (rocks, stones, wood…) carried by the water flow inside caves. In 2019, five HOBO MX2201 for water temperature were installed in three different caves (Subterráneo pool2, Pachón main and lateral pool, Sabinos pool1 and pool2). We used thin cord (type Cordelette 03 by BEAL, France) to attach loggers to natural reliefs, roughness or holes present on cave walls while preserving the stone and cave integrity. We used nodes to block the cord to prevent the probe from slipping. We attached a small rock to the end of the cable to keep the probe vertical on the water column. We took care to place the probe above the substrate to prevent mud deposition, choosing a permanent body of water (sufficiently deep in the dry season) in order to reduce the risk of air exposure.

After the success of these test installations, we followed the same procedure in the following years, while further securing the probes fixation. Where natural reliefs were inaccessible, we used an electric drill to make a hole for a dowel and we used a stainless steel anti-theft cable (for computers) to reinforce the system. In addition, large probes were protected by PVC drainpipe (with holes to facilitate water circulation) (Fig. 1B and Fig1-figure supplement 1).

### Statistics and graphical representations

Data analysis and graphs were generated under R software environment for statistical computing and some graphics (R Core Team, 2024) and with Excel. Principal component analysis (PCA) was performed using 3 variables (water temperature, pH, and conductivity) when comparing cave and surface waters, or surface running, surface still and cave waters. For between-caves and between-pools comparisons, two geomorphological variables were added (distance and depth from entrance). Biplots using the first two PCA axes (respectively named pca1 and pca2) were displayed, 95% confidence ellipses around the centroids were superimposed on the biplots. ANOVAs were conducted on the coordinates of the individuals on the first and second PCA axes. When ANOVAs were significant, post hoc tests - Tukey contrast tests - were performed.

## Acknowledgements

Since 2009, this work was supported by ANR grants (ASTYCO and BLINDTEST) and Equipe FRM grants (DEQ20150331745 and EQU202003010144) to SR, IDEV grants (2012 and 2018) to SR and Didier Casane, an ECOS-Nord Franco-Mexican exchange program grant (M15A03) to SR and Patricia Ornelas-García, a CNRS MITI grant (Mission pour les Initiatives Transverses et Interdisciplinaires) to SR. We also thank the RESAMA network, and Frédéric Sohm as Director of the previous UMS AMAGEN for financial support and for lending some equipment.

Many thanks to current and former members of the Rétaux team for help and collaborative spirit during the field expeditions, to Didier Casane for support in field work, to Patricia Ornelas-García for sharing field permits, to Jean-Louis Lacaille Múzquiz and Alejandro Durán (CONANP) for helping with access to caves located in Reservas de la Biosfera, and to Christophe Hanot for help formatting the data. We are also grateful to Stéphane Rode and Julie Rode from the NGO OSI (participative science) for technical support.

## Author’s Contributions

L.L. and S.R. designed the study; L.L. and S.P. performed the field measurement with the help of L.E. and S.R.; L.L. and S.R. analyzed the results with the help of F.R. and J.A.; SR and L.L. wrote the article.

## Supplemental Figures and Legends

**Figure 1-figure supplement 1:**
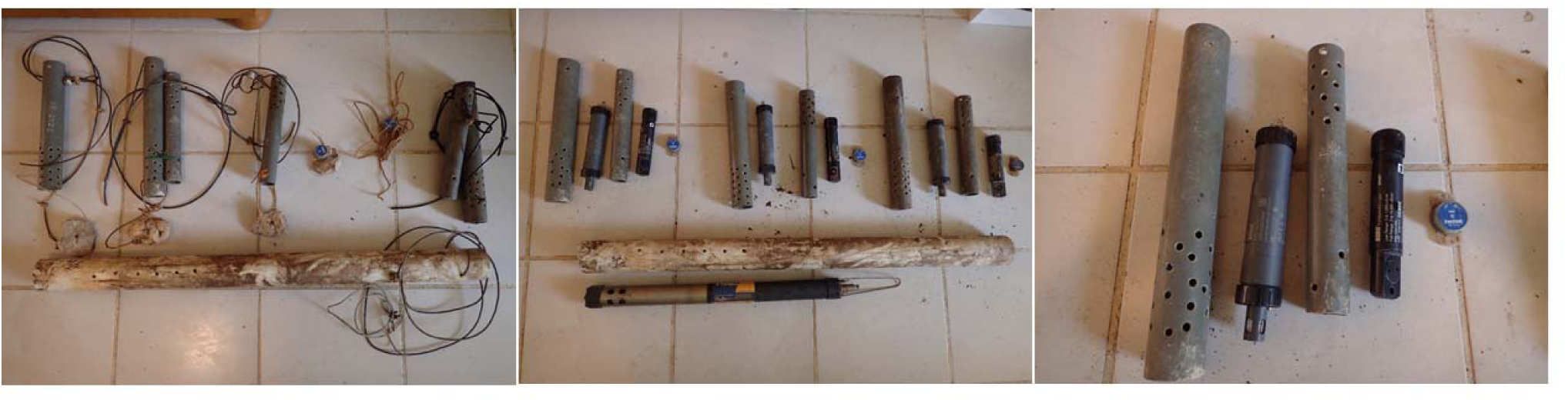
Equipment used. Photographs illustrating the state of conservation of probes (VuSitu Aquatroll600 and Hobo probes) and their fixation and protection systems, after one year spent in the natural cave environment. Photos by LL.

**Figure 2-figure supplement 1:**
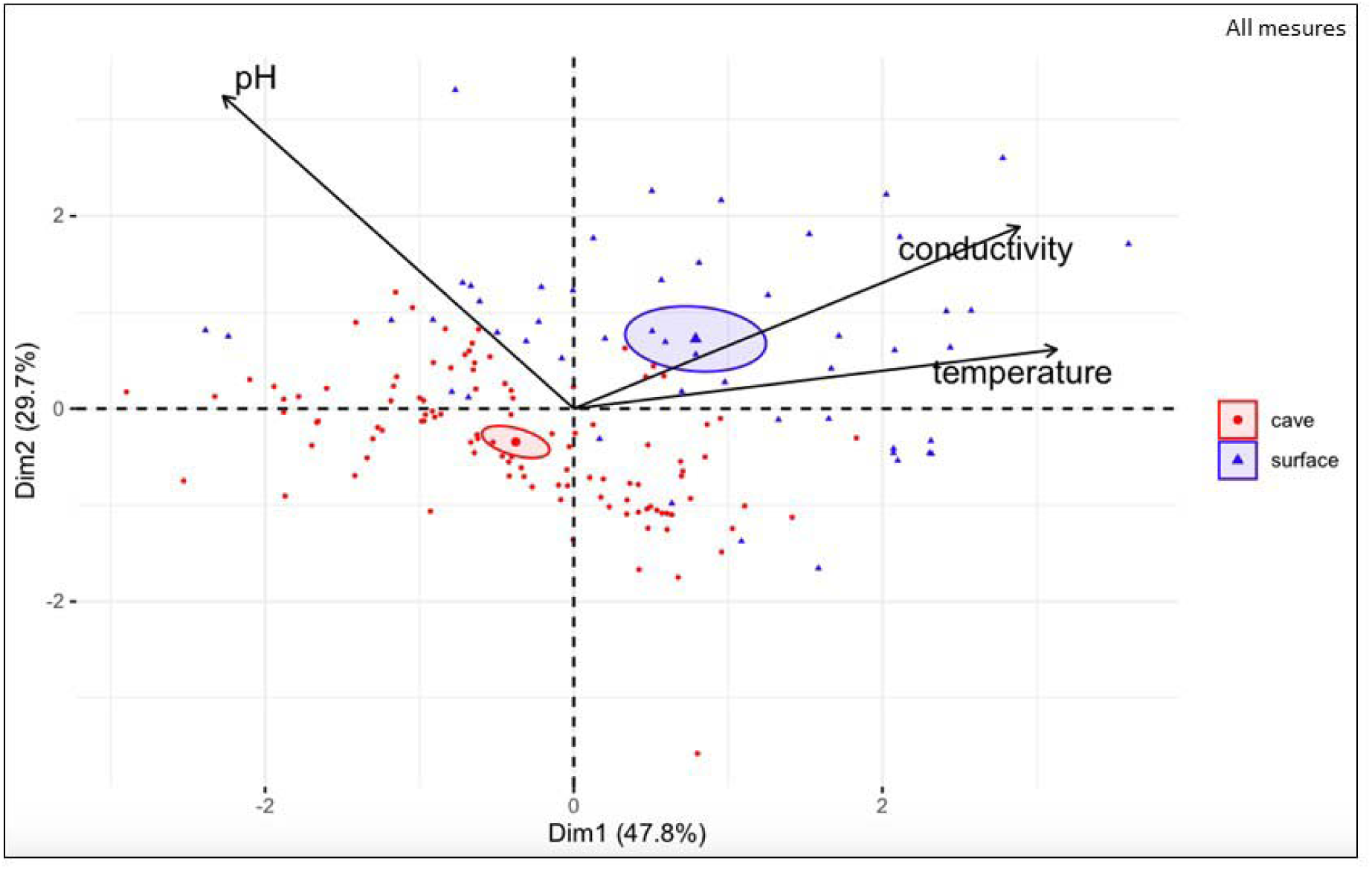
Clustering of the physico-chemical properties of rivers and caves water using PCA. This graph includes all values recorded with different instruments. Ncaves=107, Nsurface=51, N=3 variables. See Supplemental File 2, raw data. Color codes are indicated, 95% confidence ellipses are shown. Anova/pca1, pvalue=1.88e-09; Anova/pca2, pvalue=8.64e-13. The results are identical to the analysis shown in Fig. 2, where only the values recorded from Combo Hanna are shown. This strongly suggests that measures recorded with different instruments are similar and therefore reliable in our dataset.

**Figure 3-figure supplement 1:**
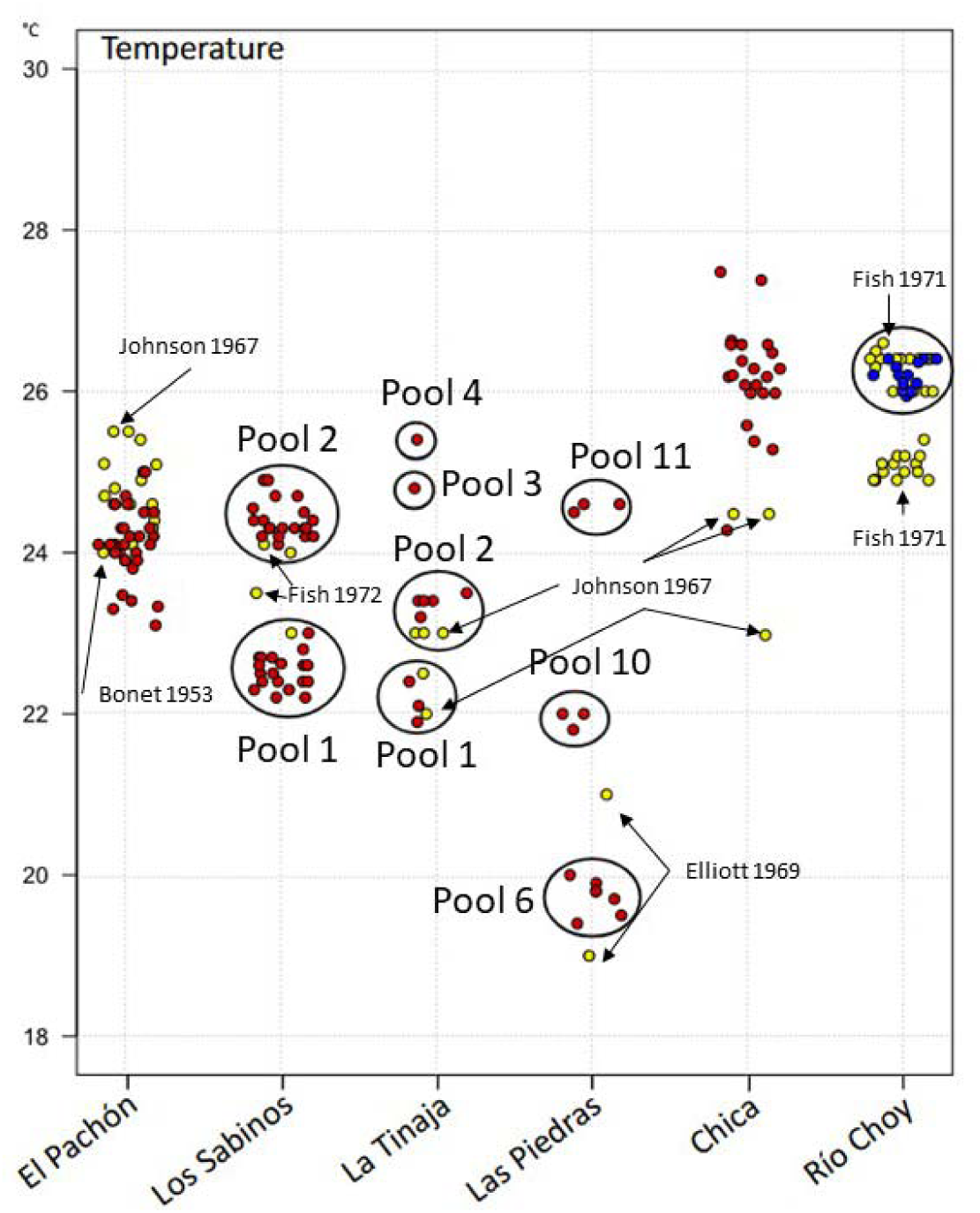
Stability of water temperature measurements over 7 decades (1953-2024) Same as Fig. 3, including also data collected for Cueva de Los Sabinos, Cueva de la Tinaja, Sótano de Las Piedras, Cueva Chica. Red dots (caves) and blue dots (river) are from our dataset. Black circles indicate cave pool numbers, known from our dataset. Yellow dots are from ancient literature, with the author name & date. However, in most cases there is no pool location information in these reports. Older measurements are in the range of ours, suggesting global stability of mean temperatures in these caves over 70 years. For the surface station Nacimiento del Río Choy (blue dots; river location #3 on Fig.1A), a black circle shows a group of ours & old data (Fish, 2004) close to 26°C. Another group of only older data is close to 25°C, also from Fish (2004). Fish showed that temperature could vary within days with flooding (e.g., between 5 June and 14 August 1971) at this location, with also massive variations in discharge, turbidity and water chemistry.

**Figure 4-figure supplement 1:**
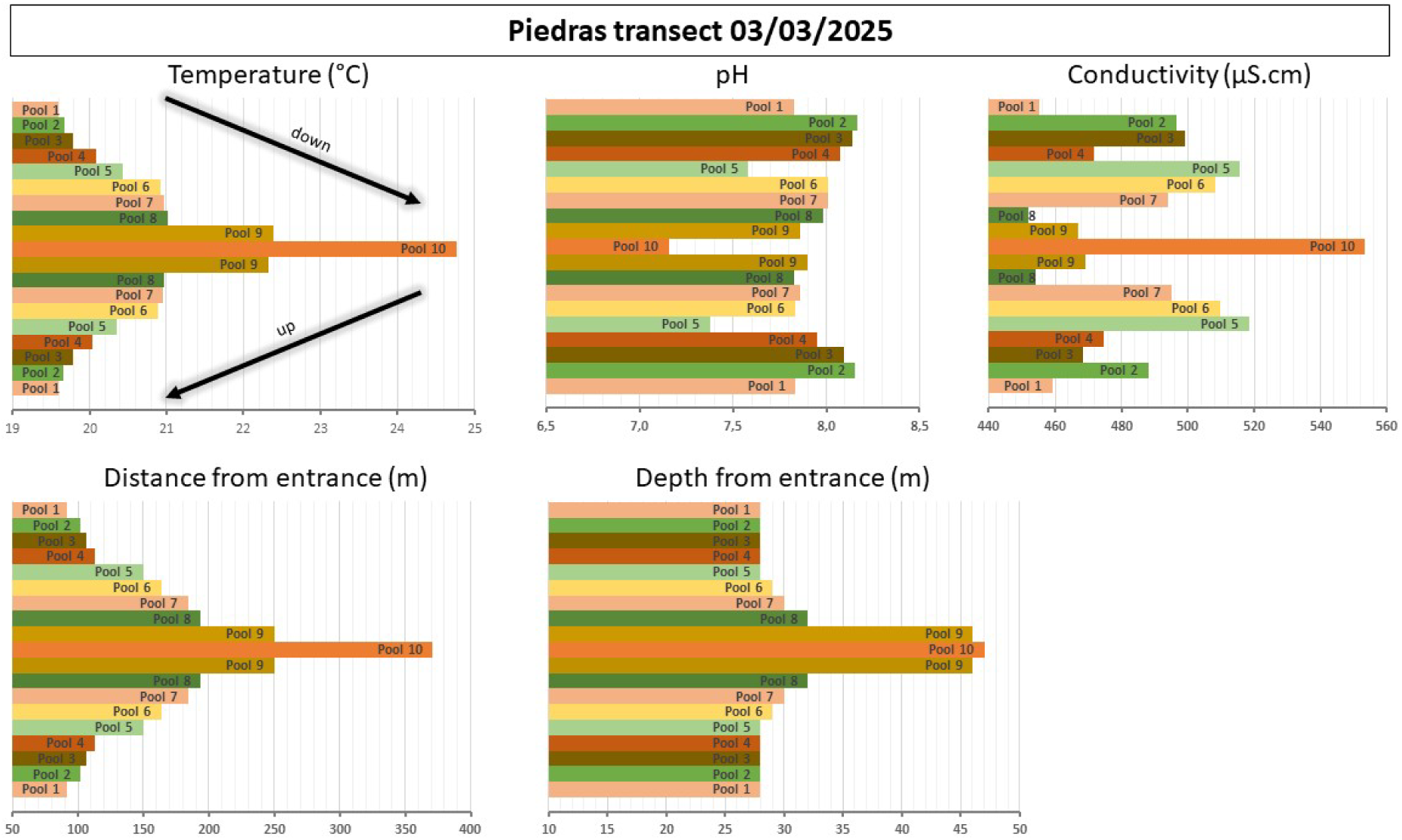
Piedras transect study, pool1 to pool10, complete dataset. Transect analysis in Piedras cave through pools 1 to 10, showing temperature, pH, conductivity, as well as distance from entrance and depth. The low-to-high temperature gradient from the first to last pool encountered seems to vary as a function of the distance from entrance, less so with depth of the pool. pH and conductivity do not show such a gradient.

**Figure 6-figure supplement 1:**
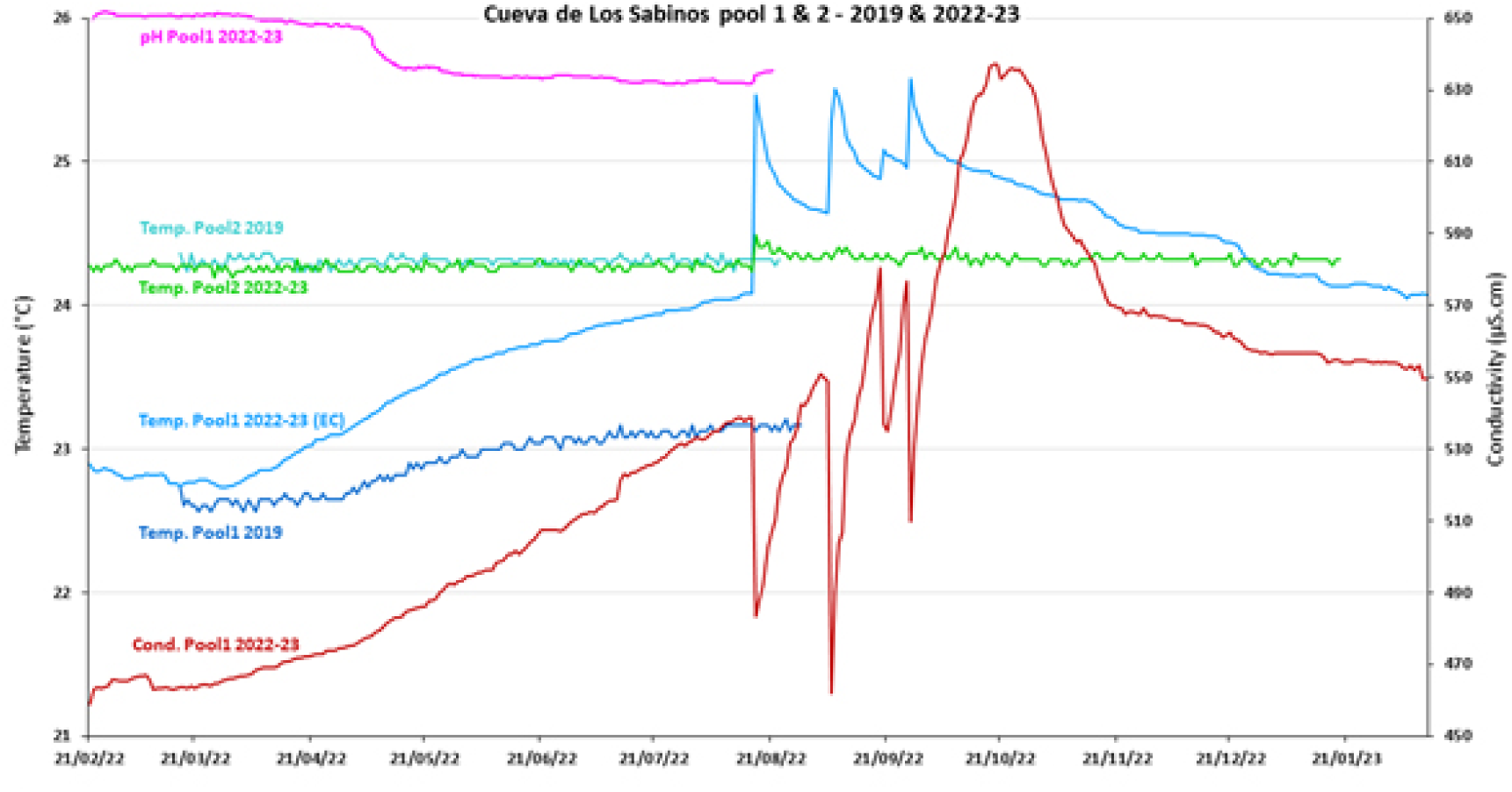
Recordings in Sabinos. The evolution of temperature, pH and conductivity, throughout the year 2019 and from February 2022 to February 2023 in Sabinos cave pools 1 and 2. Color codes for recorded parameter, pool and year of interest are indicated. The temperature in Pool1, year 2022-23, was recorded in duplicate from the conductivity probe (EC, shown) and the pH probe (not shown, for graph clarity), and they showed exactly the same pattern with a slight shift of 0.3-0.4°C all along the curve. Note that Pool2 is globally warmer than Pool1 over two years except during the rainy season; that Pool1 seems to undergo more fluctuations than Pool2; and that different parameters vary in a correlated manner in Pool1 at a daily timescale.

**Figure 8-figure supplement 1:**
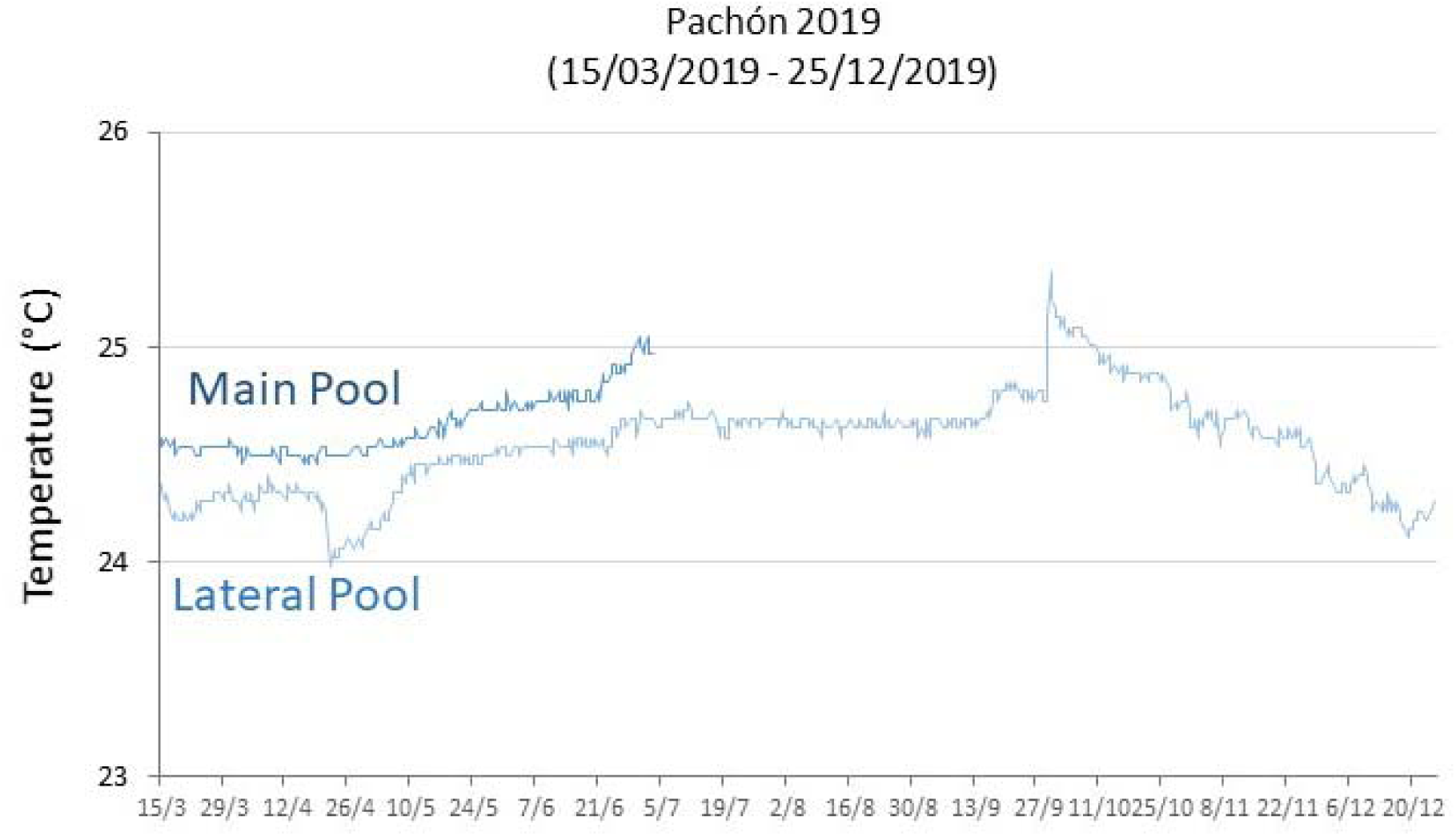
Recordings of water temperature in Pachón in 2019. Evolution of temperature from 15 March 2019 to 25 December 2019 in the main and lateral pools of the Pachón cave (Hobo probes). Recordings lasted 4 months in the main pool and 9 months in the lateral pool. Contrarily to other caves, the temperature appears very stable, varying mostly in the range between 24°C and 25°C.

