## Supplemental file 1 for "Water parameters and hydrodynamics in rivers and caves hosting *Astyanax mexicanus* populations reveal macro-, meso- and micro-habitat characteristics"

### Maps of caves studied and sampling locations

### Cueva de El Pachón

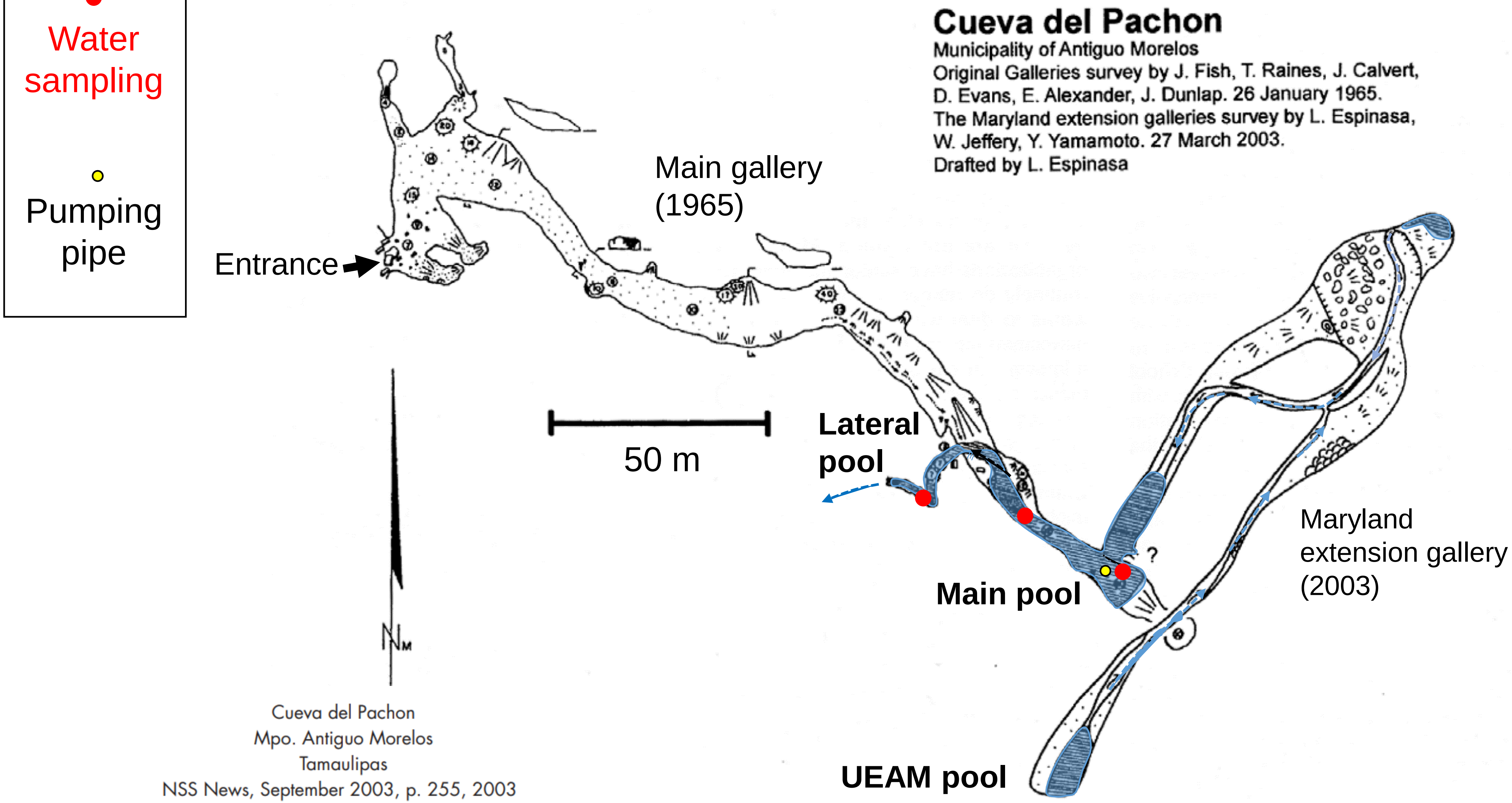


Cueva del Pachon
Mpo. Antiguo Morelos
Tamaulipas
NSS News, September 2003, p. 255, 2003
Astyanax International Meeting 2009 program and abstracts, p. 16, 2009

Water sampling

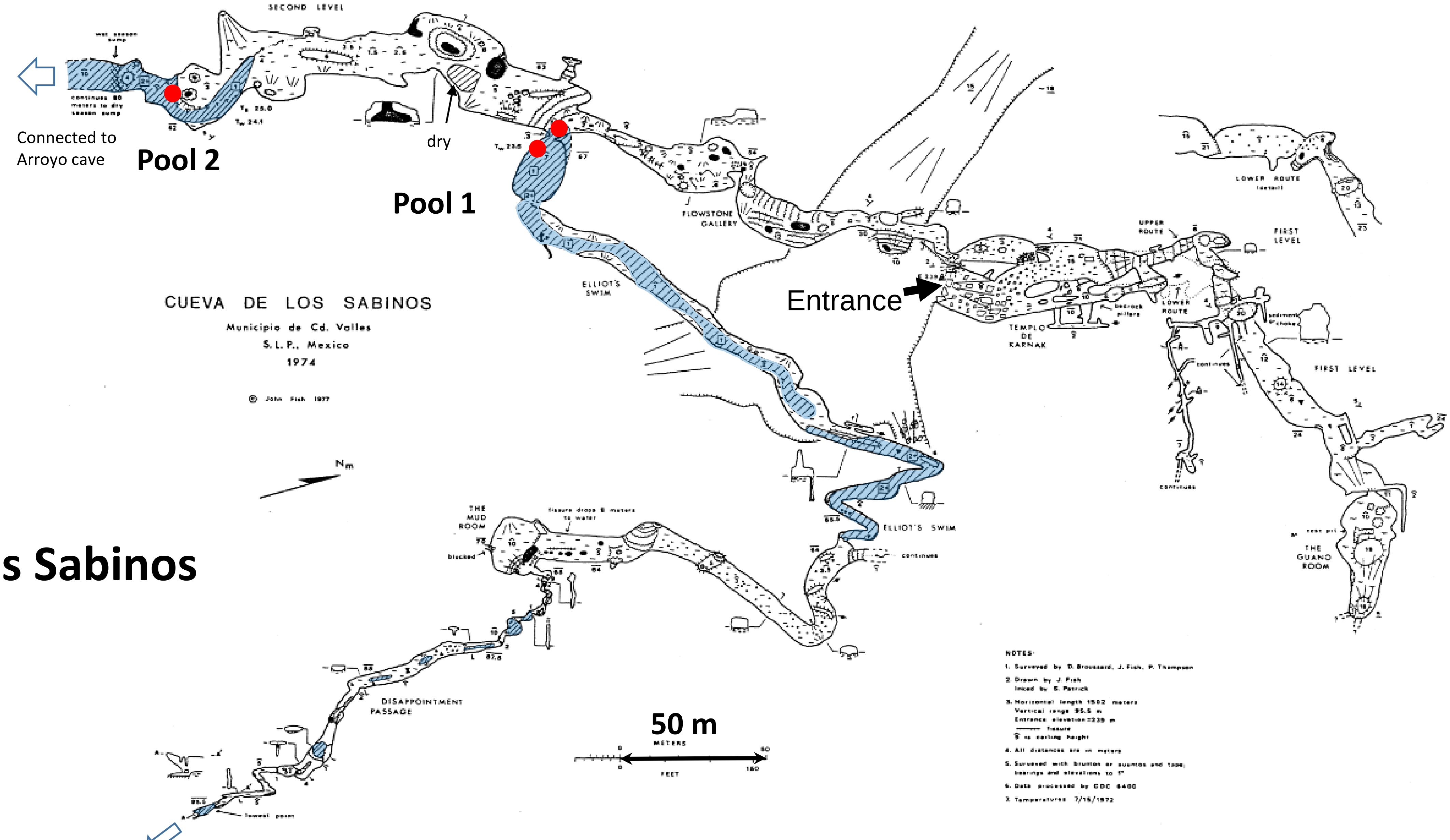


Probably connected to Arroyo cave

### Cueva de Los Sabinos

Cueva de Los Sabinos
Mpo. Valles
San Luis Potosí
Karst Hydrology of the Sierra de El Ebra, Mexico, p. 117
Canadian Caver, v. 6, no. 1, p.15, 1974

### Sótano de las Piedras

Water sampling

Brunton compass & tape surveys by William R. Elliott, Jim McIntire & Don Broussard, 17 July 1969; William R. Elliott, Robert Hemperly, John Prentice & Carmen Soileau, 19-20 May 1974. Entrance elevation 145 m, depth 52 m, horizontal length 434 m, extent 468 m. Redrawn by Elliott, 2014.

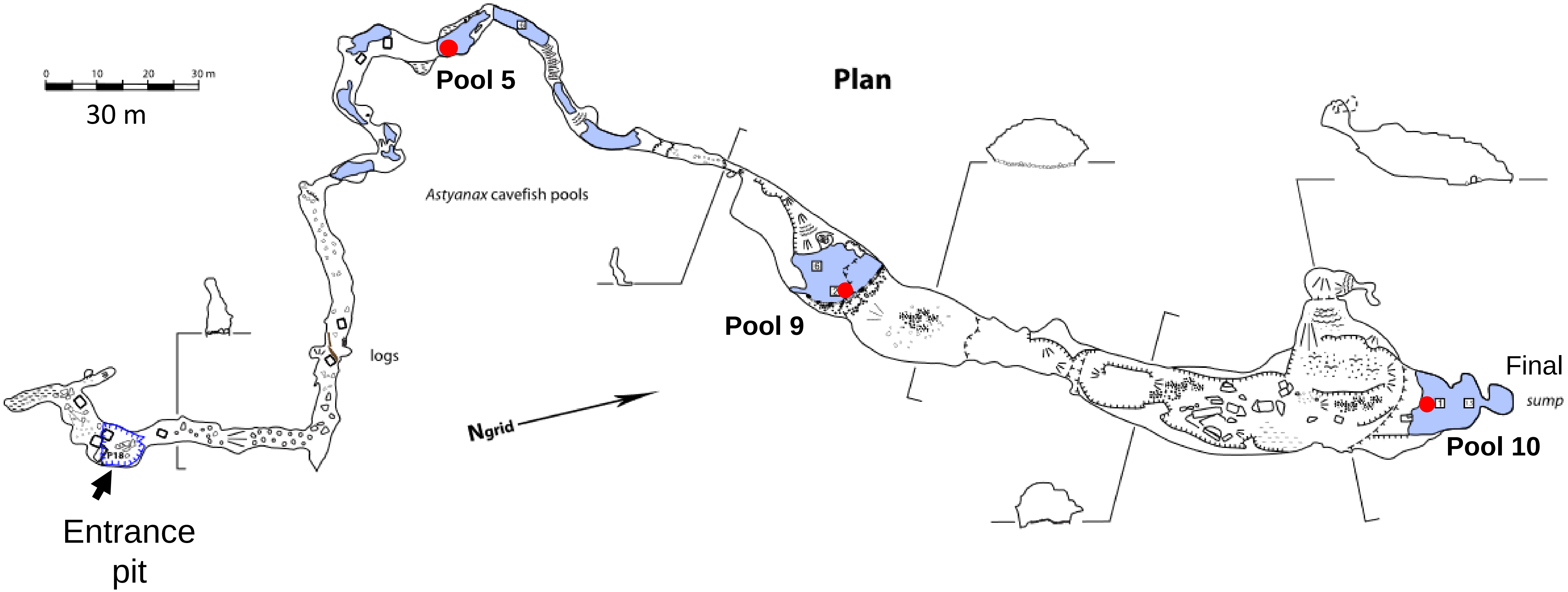

Water sampling

### Sótano de la Tinaja

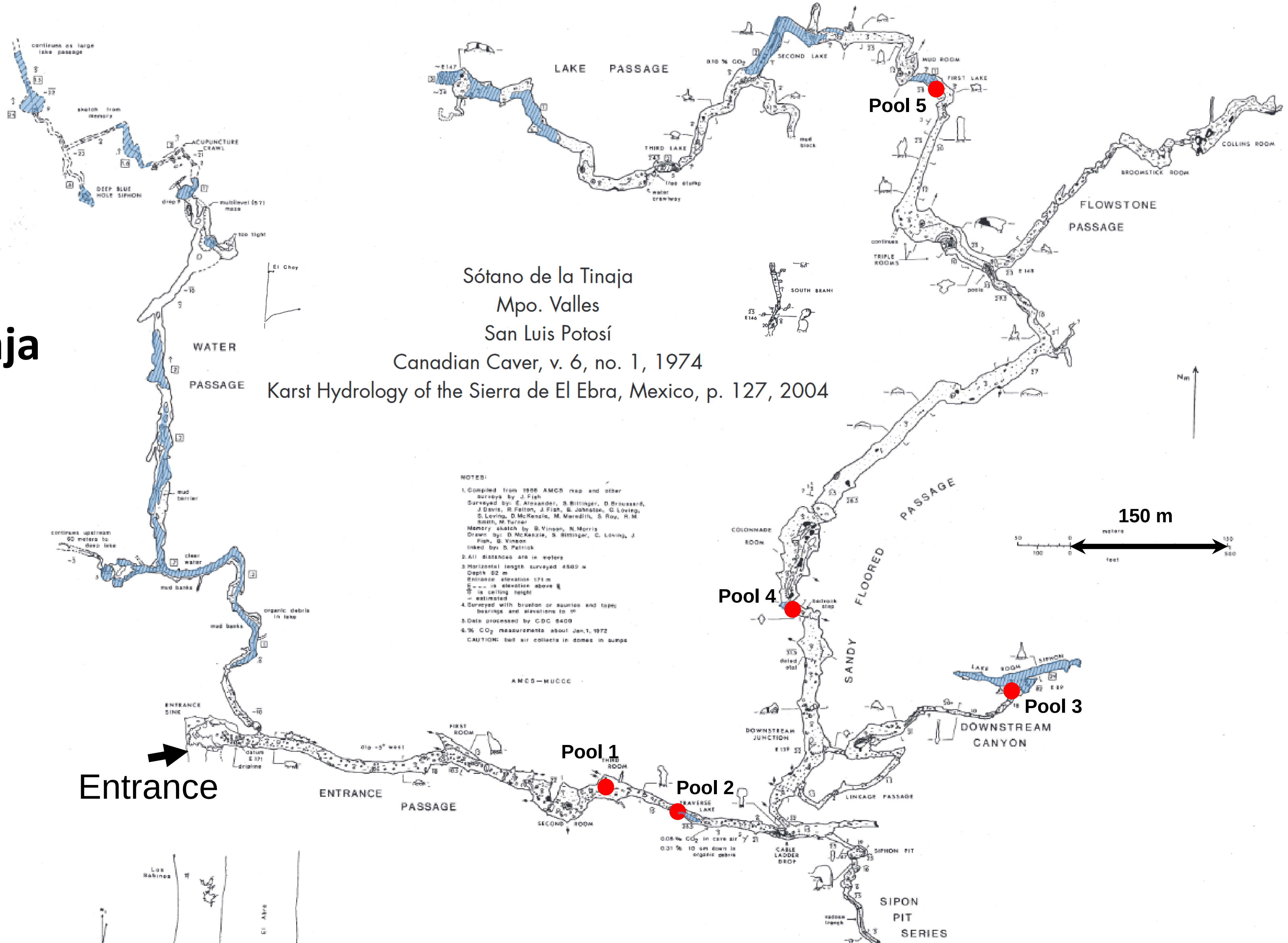

### Cueva Chica

●
Water sampling

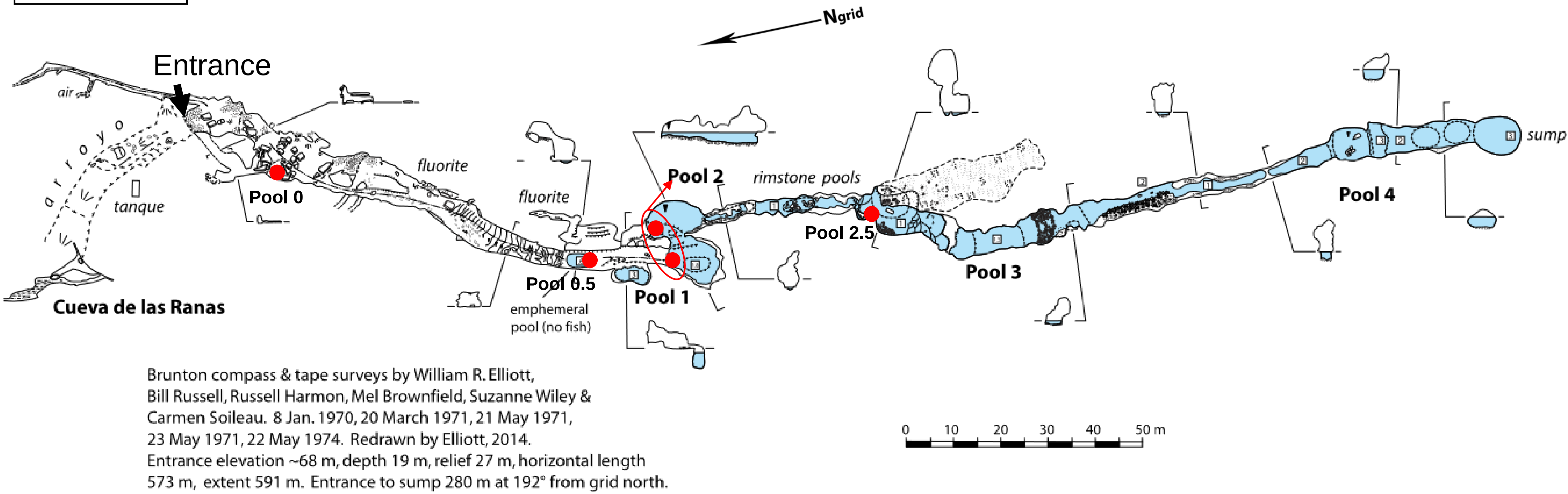
Entrance
Ngrid
air
arroyo
tanque
Cueva de las Ranas
Pool 0
fluorite
fluorite
Pool 0.5
emphemeral
pool (no fish)
Pool 1
Pool 2
rimstone pools
Pool 2.5
Pool 3
Pool 4
sump
0 10 20 30 40 50 m
Brunton compass & tape surveys by William R. Elliott,
Bill Russell, Russell Harmon, Mel Brownfield, Suzanne Wiley &
Carmen Soileau. 8 Jan. 1970, 20 March 1971, 21 May 1971,
23 May 1971, 22 May 1974. Redrawn by Elliott, 2014.
Entrance elevation ~68 m, depth 19 m, relief 27 m, horizontal length
573 m, extent 591 m. Entrance to sump 280 m at 192° from grid north.

### Cueva del Río Subterráneo

Water sampling

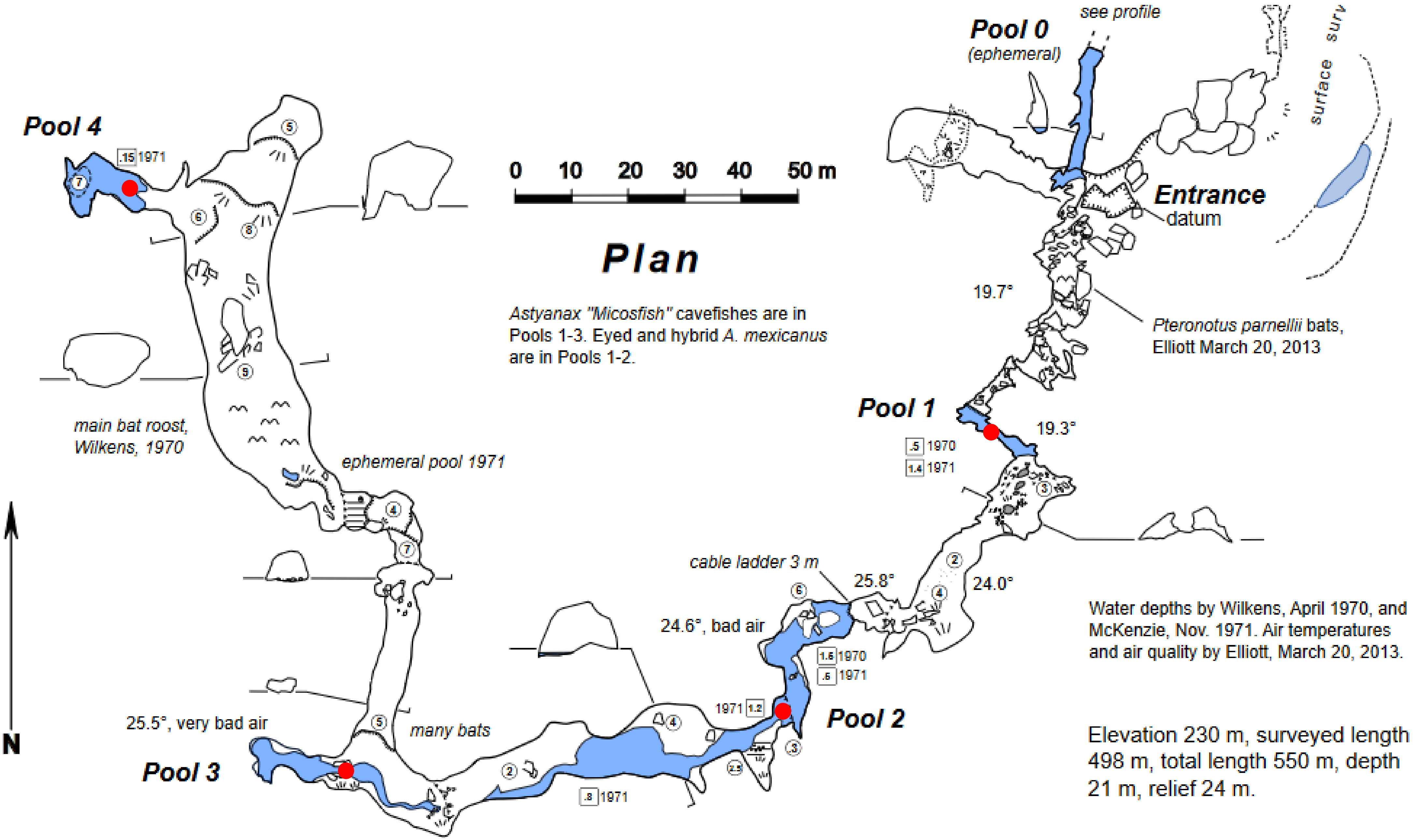
